# The world’s hotspot of linguistic and biocultural diversity under threat

**DOI:** 10.1101/2021.04.12.439439

**Authors:** Alfred Kik, Martin Adamec, Alexandra Y. Aikhenvald, Jarmila Bajzekova, Nigel Baro, Claire Bowern, Robert K. Colwell, Pavel Drozd, Pavel Duda, Sentiko Ibalim, Leonardo R. Jorge, Jane Mogina, Ben Ruli, Katerina Sam, Hannah Sarvasy, Simon Saulei, George D. Weiblen, Jan Zrzavy, Vojtech Novotny

## Abstract

Papua New Guinea is home to >10% of the world’s languages and rich and varied biocultural knowledge, but the future of this diversity remains unclear. We measured language skills of 6,190 students speaking 392 languages (5.5% of the global total) and modelled their future trends, using individual-level variables characterizing family language use, socio-economic conditions, student’s skills, and language traits. This approach showed that only 58% of the students, compared to 91% of their parents, were fluent in indigenous languages, while the trends in key drivers of language skills (language use at home, proportion of mixed-language families, urbanization, students’ traditional skills) predicted accelerating decline of fluency, to an estimated 26% in the next generation of students. Ethnobiological knowledge declined in close parallel with language skills. Varied medicinal plant uses known to the students speaking indigenous languages are replaced by a few, mostly non-native species for the students speaking English or Tok Pisin, the national lingua franca. Most (88%) students want to teach indigenous language to their children. While crucial for keeping languages alive, this intention faces powerful external pressures as key factors (education, cash economy, road networks, urbanization) associated with language attrition are valued in contemporary society.

**Significance Statement:** Around the world, more than 7,000 languages are spoken, most of them by small populations of speakers in the tropics. Globalization puts small languages at a disadvantage, but our understanding of the drivers and rate of language loss remains incomplete. When we tested key factors causing language attrition among Papua New Guinean students speaking 392 different indigenous languages, we found an unexpectedly rapid decline in their language skills compared to their parents and predicted further acceleration of language loss in the next generation. Language attrition was accompanied by decline in the traditional knowledge of nature among the students, pointing to an uncertain future for languages and biocultural knowledge in the most linguistically diverse place on Earth.

---

When evaluated against a common set of extinction risk criteria, the world’s ∼7,000 extant languages (1) are even more threatened than its biological diversity (2). Orally transmitted cultural knowledge may be threatened by similar forces (3, 4). Language population sizes approximate a log-normal distribution (5), such that the majority of languages have relatively few speakers (1). Nearly half of the world’s languages are considered endangered (1, 6). Language extinction is accelerating, with 30% of recorded extinctions having occurred since 1960 (6). Language vulnerability to extinction depends on speakers’ attitudes toward their languages, as well as on socio-economic factors (7). However, quantitative evidence on the relative impact of individual drivers of language endangerment is almost non-existent (8, 9), making it impossible to understand and predict language attrition. Further, language skills and ethnobiological knowledge are rarely examined in relation to socio-economic variables for individual speakers, as required for mechanistic understanding of language attrition and loss of ethnobiological knowledge (10, 11, 12). The present study uses a novel modelling approach to assess multiple drivers of language attrition and ethnobiological knowledge loss based on extensive data for individual speakers to predict future trends in a global hotspot of linguistic and cultural diversity.

Papua New Guinea (PNG) is the world’s most linguistically diverse nation, where ∼9 million people speak ∼840 languages (5, 13). PNG’s languages are highly diverse, classified into at least 33 families (14). Until recently, these languages enjoyed widespread vitality, due to the absence of a dominant language in the region, stable small-scale multilingualism (15), and focus on language as a marker of group identity (7, 16). New Guinea is also the world’s most floristically diverse island (17), comprising approximately 5% of the world’s biodiversity (18). Throughout PNG, numerous indigenous communities have explored, systematized, used and managed the extraordinary biodiversity in their natural environment, thus generating extensive biocultural knowledge of local ecosystems (19, 20, 21). The traditional environmental knowledge of indigenous communities is in decline world-wide, in response to the forces of cultural and economic globalization (22). Only 20% of PNG ethnolinguistic groups have any of their traditional plant uses recorded in the literature, and detailed information (>100 plant use records) exists for only 2.5% of groups (3). Likewise, the contemporary status on this knowledge remains poorly documented.

At present, 32% of indigenous languages in PNG are considered endangered (1), largely due to their replacement by Tok Pisin (an English-based creole, and PNG’s major lingua franca) or English (the language of formal education) (23). However, the true status of the country’s languages cannot be assessed in the absence of a national linguistic survey (24). This study presents such a survey and examines the present status and future dynamics of language and biocultural knowledge loss.

## Results and Discussion

### Language skills drivers

We used questionnaires that compiled information on socio-economic background and self-reported language fluency for 6,190 secondary school students, followed by tests of their language skills and ethnobiological knowledge. This survey captured 392 languages (46% of languages spoken in PNG and 5.5% worldwide), including 110 languages with ≥10 respondents (Fig. 1, Dataset S1). We have uncovered a dramatic decline in the language skills in a single generation. While 90.8% of students’ parents reportedly speak an indigenous language fluently and only 0.3% of them have no indigenous language skills, just 57.7% of students consider themselves fluent in an indigenous language, whereas 2.0% of students reported a complete lack of indigenous language (Fig. 2A). The 110 languages with ≥10 respondents lost, on average, 40±2.1% (±s.e.) of fluent speakers in the contemporary generation, from parents to the secondary school students we studied (Fig. 2B). The parent-student comparison suggests that language attrition is a recent phenomenon and thus not a direct consequence of the colonial past of PNG (until 1975), but rather a result of economic and social development of a country undergoing globalization.

**Figure 1.**
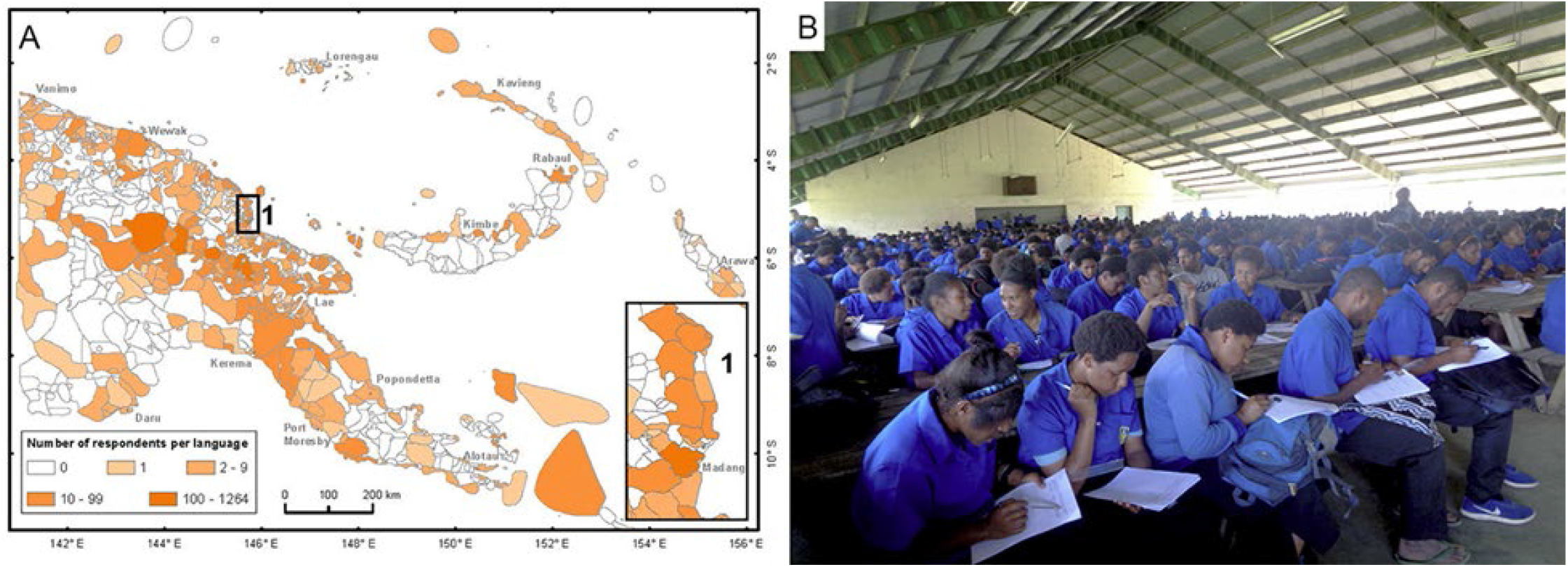
Languages studied in Papua New Guinea. (A) Language map (*1*) with the number of students surveyed, (B) Survey of 486 students, speaking 37 indigenous languages, at the Mt. Hagen Secondary School.

**Figure 2.**
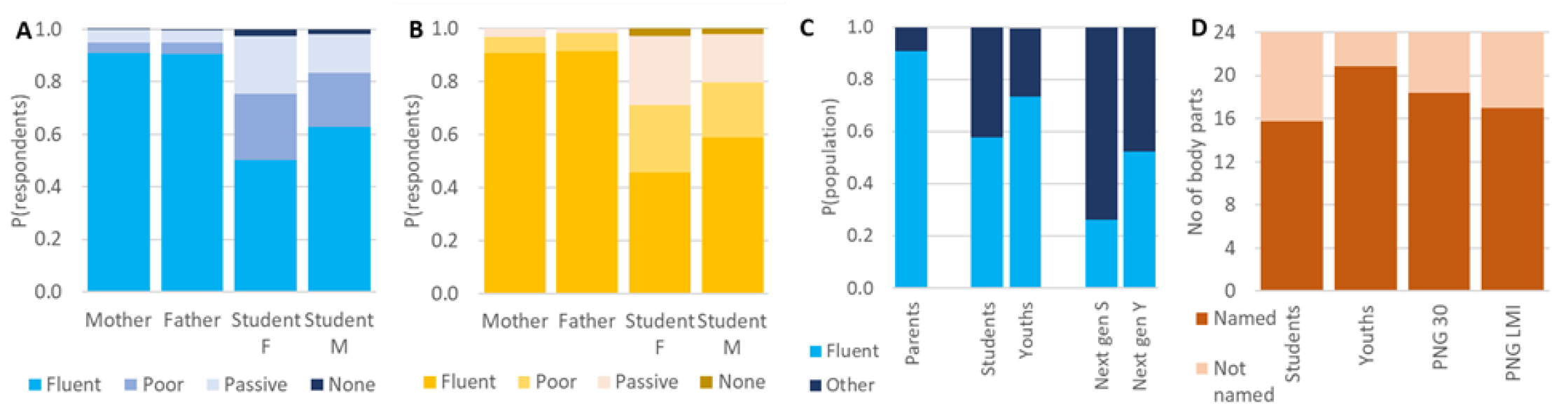
Indigenous language skills in present and future PNG populations. (A) Language skills (L2) of 6,190 students (female and male) and their parents, (B) Mean language skills (L2) for 110 well-sampled languages (N ≥ 10 students per language), (C) The proportion of fluent speakers among parents and students, extrapolated to the entire 18-20 year-old cohort in PNG (Youths), and to the next generation of students (Next gen S) and all 18-20-year-olds (Next gen Y), (D) Language skills (L1) of the students, predictions from models characterizing the 18-20-year-olds in PNG at present (Youths) and in 30 years (PNG 30), and language skills assuming that PNG will come to match the mean socio-economic parameters of Lower-Middle Income countries (PNG LMI). Language skills were quantified as the number of body parts (from the total of 24) correctly named from photographs (L1), or by assessment by respondents for themselves and their parents on four-point scale: 0 - no language skills, 1 - passive understanding, 2 - speaking but poorly, 3 - fluent use (L2).

We tested a set of factors characterizing student’s life skills, family language use, socio-economic conditions, and language traits that potentially affect language skills (7) (Fig. 3, SI Appendix, Fig. S1 and Table S1). The language used at home was the most important predictor of language skills. Indigenous languages, used in 30% of all families, competed with Tok Pisin and English, used respectively in 66% and 4% of families. More interestingly, home language use was also strongly impacted by mixed-language family background, the second most important predictor of language skills. The effect of mixed-language family remained large even after taking into account its effect on home language use, since only 16% of mixed-language families used an indigenous language at home, compared to 38% of same-language families.

**Figure 3.**
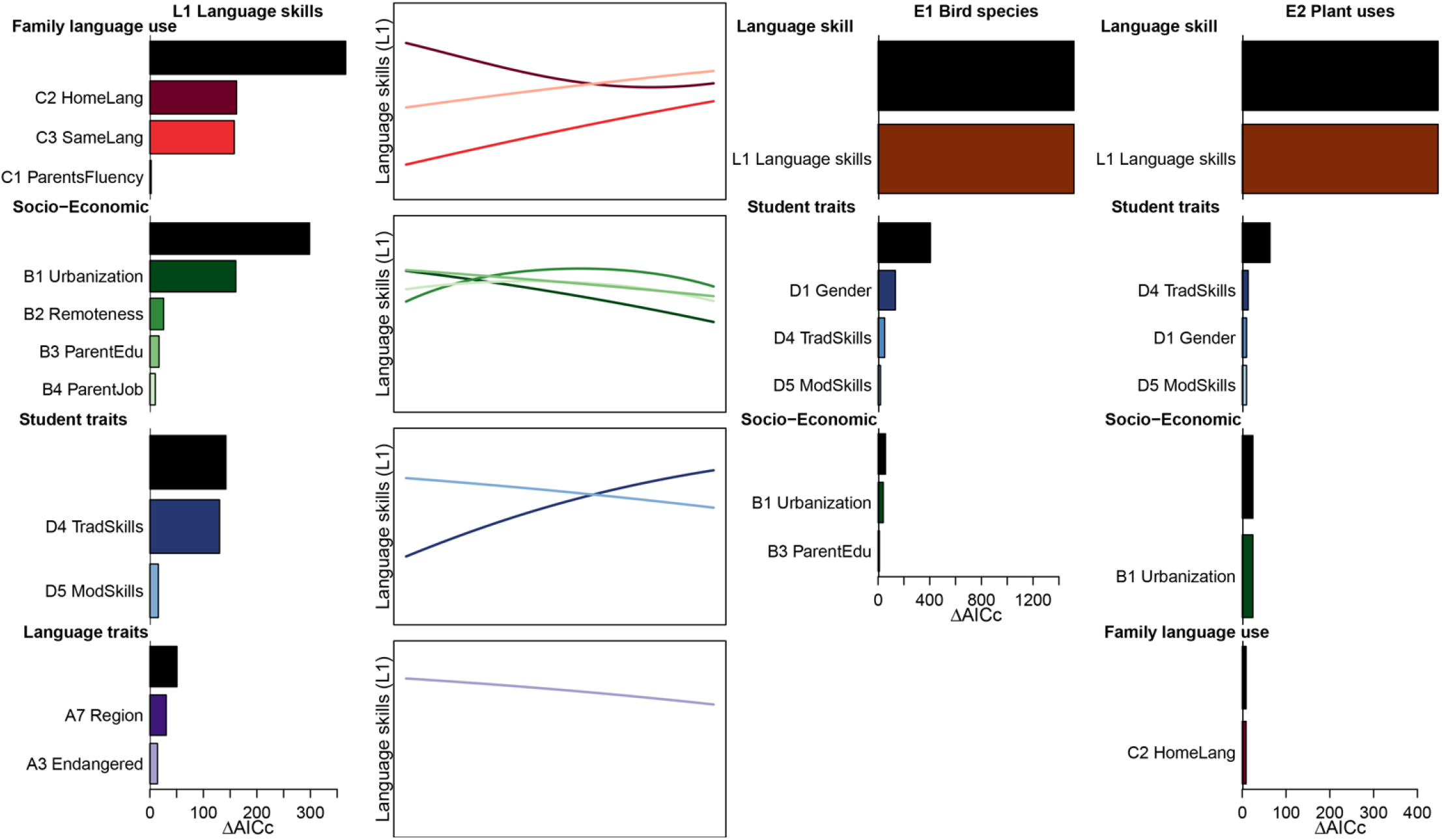
Effects of language and socio-economic factors on indigenous language skills and ethnobiological knowledge. Generalized linear mixed models (GLMM) describe variability in student language skills (L1), and in knowledge of bird species (E1) and traditional plant uses (E2). The language skills model incorporated 12 fixed variables divided into four classes: (A) Language traits: endangerment (A3), geographic regions (A4); (B) Socio-economic traits: birthplace urbanization (B1) and remoteness (B2), parents’ education (B3) and employment (B4); (C) Family language use: parents’ language skills (C1), home language use (C2), and whether parents speak the same first language (C3); (D) Student traits: traditional skills (hunting, fishing, growing food, house building, medicinal plants) (D4), and contemporary technical skills (mobile phone and computer use) (D5). The variables were selected within each class (SI Appendix, Table S2) before being included in a global model (SI Appendix, Table S3). The bars show the AIC improvement due to the addition of each group (black) and each variable within each group into a model that includes all other variables, quantifying the marginal effect of each class/variable. The line plots show the shape of the effect of each variable across its range (except categorical A4), while keeping the other variables constant. Only significant (P<0.05) variables are shown. The models describing variability in student knowledge of bird species (E1) and traditional plant uses (E2) used language skills (L1) and three classes of explanatory variables (Family language use, Socio-economic traits, and Student traits including D1 – gender) (SI Appendix, Tables S5, S6). L1 is defined in Fig. 2, other variables in Materials and Methods.

The small-scale multilingualism that was historically widespread in PNG and continues in rural parts of the country (25, 26) does not lead to language attrition (27). However, modern urban mixing, with communication in Tok Pisin or English, is different (28). Presently, 37% of the surveyed students grew up in mixed-language families. The secondary schools we surveyed are a favorable environment for language mixing, attended by students speaking 17 – 124 languages per school (SI Appendix, Table S2). Only 35% of the students speak the same indigenous language as their best friend, which is not very different from the 23% of students expected to do so if friendships were formed randomly with respect to the languages spoken by students. This pattern indicates a potential for further increase in non-traditional mixed-language marriages of these students.

Urbanization, another important factor correlated with language skills, often interrupts contacts between generations, crucial for language transfer (7, 10). Urbanization in PNG has been kept low (87% of the population is rural) (29) by customary land ownership (92% of families in our study owned land), since urban dwellers could lose their land rights to relatives who continue to live on their land in villages (30). Urban environment had a strong negative impact on language skills among the 35% of students growing up in towns and cities, compared to growing up in a rural setting, particularly in a remote village.

Parents’ education and employment had only small effects on language skills once the related factors of urbanization and home language use were accounted for (Fig. 3). Students whose parents had salaried employment had lower language skills compared to those with parents growing cash crops or food for subsistence. The statistical importance of parents’ language skills was low, since almost all were fluent in an indigenous language. Indigenous language skills were positively correlated with a student’s reported traditional skills (hunting, fishing, farming, house building, and medicinal plant use), and negatively with contemporary technical skills (mobile phone and computer use). The individual differences in student skills thus remain important within both rural and urban environments, apart from a large decline in traditional skills and improvement in contemporary technical skills associated with transition from rural to urban lifestyle. We did not survey changes in traditional skills between students and their parents, but it is likely that good farming skills, in particular, are almost universal among the parents, compared to 68% for the students. Interestingly, the student’s English skills and mathematical skills had no effect on language skills, showing the limited direct effect of formal school education compared to life-style changes. Finally, language skills did not differ between female and male students.

The EGIDS (31) language endangerment classification, based on inter-generation transfer of languages and social domains of their use, is a significant predictor of language skills. Our study thus validates this endangerment parameter. Unlike some other measures of endangerment (2, 8), EGIDS does not consider the number of speakers of a language. We tested language size separately and found it had no significant effect on language skills; this finding bodes well for the survival prospects of numerous small languages in PNG (in 2000, the median language had only 1,201 speakers, Dataset S1).

In PNG, 87% of languages have a writing system, but only 15% of those languages have even a limited dictionary (1). Literature is thought to promote language vitality (31), but the existence of a Bible translation, typically the only written text in indigenous languages of PNG, did not improve language skills for the 84% of students who speak indigenous languages with Bible translations. This result could reflect the fact that only a third of Bible translations are extensively used (32). The students’ language skills also differ across geographic regions of PNG, probably reflecting regional differences in environmental or socio-economic factors not directly captured by the analysis.

While many of the language attrition drivers we detected have been documented previously (8, 10, 33), our analysis quantified their relative importance and revealed that multiple factors, even when correlated, have significant, statistically independent effects. For instance, urban lifestyle was correlated with better education and salaried employment of parents, and with low traditional and high contemporary technical skills of students, but all these variables remained significant, independent predictors of language skills (SI Appendix, Fig. S2).

### Future trends in language skills

Only 15% of young people in PNG attend secondary school (34). They tend to come from towns and cities (35% in our sample vs. 13% in the general population), have educated parents (17% with tertiary education vs. 5% country-wide in the age cohort 45-54 years), and rely less on subsistence agriculture (31% vs. 57% in the general population) (34) (SI Appendix, Table S1). Rural families can often afford education beyond primary level for only one or a minority of their children.

Considering these selection biases, we estimated indigenous language fluency for the entire 18-20-year-old cohort of PNG, using country-wide values for urbanization, parents’ education, and parents’ employment as independent variables. We also used these variables to estimate the country-wide proportion of linguistically mixed families and the proportion of households using any indigenous language. Our model estimated that 73.5% of the 18-20-year-olds in PNG are fluent in an indigenous language - a higher proportion than among secondary students but representing a significant decline from their parents, of whom >90% are likely fluent in at least one indigenous language (Fig. 2C). While the share of fluent speakers decreased dramatically from parents to their children, the PNG population almost doubled during the same period, from 4.62 million in 1990 to 8.95 million at present (29). It is predicted to grow further, to 27 million in 2100 (35). The absolute number of fluent speakers thus probably increased in the past 30 years for most indigenous languages in PNG and may continue to grow in the future, whilst representing a rapidly diminishing share of the total population. Such an increasingly minority position may be detrimental for the survival of indigenous languages, irrespective of the number of speakers.

We used extrapolated values of language skills drivers to model the situation for students and all 18-20-year-olds in the next generation. Unlike most other countries, PNG is predicted to remain predominantly (76%) rural in 2050 (36). Higher mobility, including travel for education and employment, will likely lead to an increase in the already high proportion (37%) of linguistically mixed families; a hypothetical random selection of partners would result in 99% of mixed families, based on our population size estimates for PNG languages (Dataset S1). We used the proportion of students whose best friend speaks a different first language (65%) as a proxy for the future share of mixed-language families. The share of the population with secondary or tertiary education is expected to increase from 19% to 31% by 2050 (34), but the proportion of the population with salaried employment was modelled as constant (31%), since there has not been a definitive trend over the past 30 years (37).

Our model predicted that the current students’ 58% fluency in indigenous languages will shrink to 26% for the next generation. Further, we estimated 52% fluent indigenous language speakers in the entire 18-20-year-old cohort of the next generation in PNG (Fig. 2C). We also modelled the scenario of PNG converging to the mean socio-economic parameters for Lower-Middle Income Countries, and this model predicted even greater attrition in language skills in the general population (Fig. 2D).

### Ethnobiological knowledge in decline

We tested the knowledge of indigenous bird species and traditional uses of plants as two important components of biocultural knowledge (20). The knowledge of both bird species and plant uses was closely predicted by indigenous language skills, and therefore in decline (Fig. 3, SI Appendix, Fig. S3). This result was expected as most indigenous plant and animal names lack established translations into Tok Pisin or English and scientific species identifications (19). The continued maintenance of traditional knowledge in the face of severe language loss is rare, and this knowledge may be lost or restructured even when the indigenous language remains healthy (38, 39). Language shift, together with formal education, transition to a market economy, new technologies, urbanization, interethnic contact, habitat degradation, modern health care, religious belief, change in values, and modern media have been identified as global drivers of decline in ethnobiological knowledge, and its replacement or fusion with new information from external sources (22, 38, 39).

Male students knew birds better than female students, probably because the knowledge of birds was correlated with hunting skills, which were better developed in male students. Several other student and socio-economic traits were correlated with ethnobiological knowledge, but their importance was low (Fig. 3). The close correlation between language skills and ethnobiological knowledge may result partly from the focus of our ethnobiology tests on naming species. However, the ability to recognize and name species is a prerequisite for acquiring deeper ecological and cultural knowledge of plants and animals, as we have observed when training para-ecologists, who use their traditional knowledge of the natural world to build modern research skills (40).

Student traits, including traditional skills, and socio-economic traits, particularly urbanization, were the best predictors of ethnobiological knowledge when language skill itself was not used as an explanatory variable (SI Appendix, Fig. S4). The intricate details of biology are often learned during teenage years spent in rainforests (41), an option no longer available to many students growing up in towns or leaving villages for boarding schools. Even the iconic and culturally important cassowary (*Casuarius* spp.) (42) could be named in an indigenous language by only 64% of respondents.

The students were asked to list up to 10 plant species with their traditional uses, in indigenous languages; when they did not know any, they used Tok Pisin or English names. The majority of the plant uses reported in indigenous languages and in Tok Pisin/English were medicinal, but the proportion was greater in Tok Pisin/English responses, where 80% were medicinal, versus just 53% for plants reported in indigenous languages. Although medicinal use is often one of the most salient across cultures (20), plants reported in indigenous languages had a wide range of reported uses, including sorcery, house building, and ceremonies (Fig. 4A) (43). Further, the Tok Pisin/English medicinal uses were dominated by merely 10 plant species, only two of them (*Laportea* sp. and *Morinda citrifolia*) native to PNG (Fig. 4B). *Laportea* is widely distributed and used across PNG, while *Morinda* is a lowland species that has become commercialized throughout the Pacific (44). Students with poor indigenous language skills thus showed severely reduced traditional medicinal knowledge, replaced by an impoverished, highly “globalized” knowledge pertaining to a few, mostly non-native plant species (e.g., *Carica papaya, Citrus* spp., *Aloe vera*).

**Figure 4.**
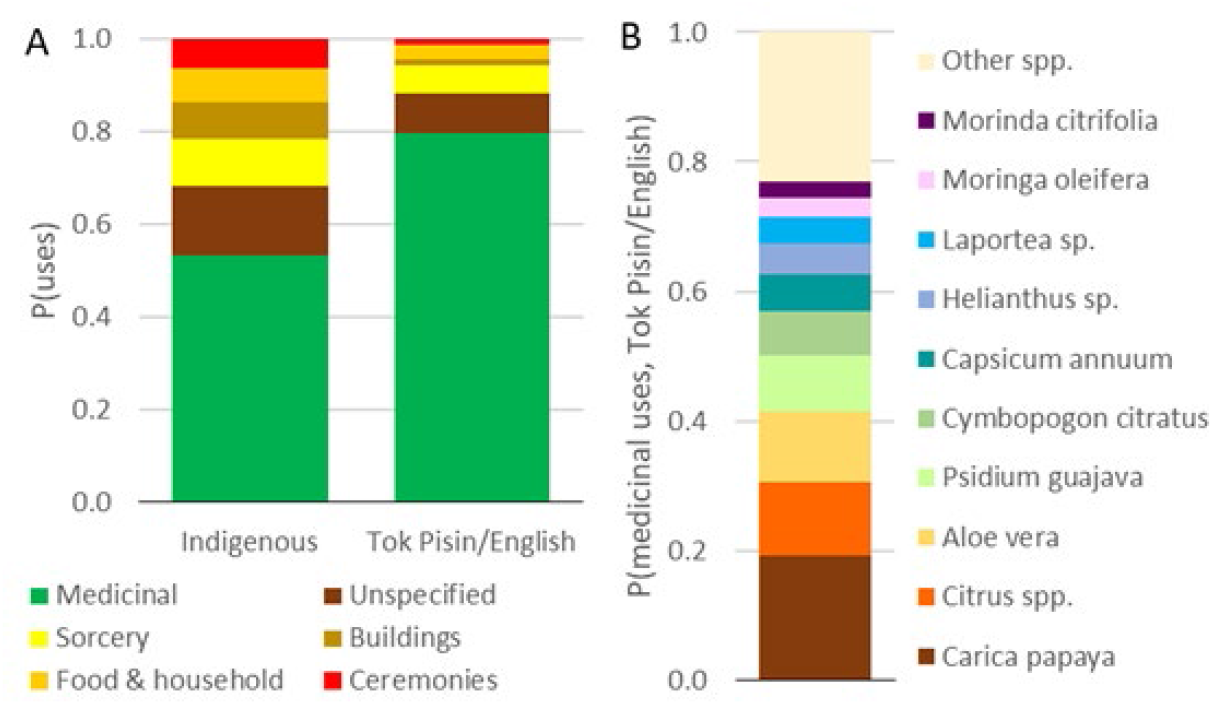
Language skills and ethnobiological knowledge. (A) indigenous plant use by categories (Hill’s diversity ^1^D = 4.15 for indigenous and 2.18 for Tok Pisin/English uses); (B) Ten most common plant species listed in Tok Pisin/English medicinal uses. The respondents were asked to freely list up to 10 plant species with their indigenous names and traditional uses (E2). They provided 21,829 responses in indigenous languages (i.e. 35% of the maximum of 10 uses x 6,190 respondents), and an additional 5,458 responses in Tok Pisin/English when they could not name any plant in an indigenous language.

## Conclusions

We have shown that the drivers of language loss documented for communities around the world (45) are, to variable extents, at play in the world’s most linguistically diverse nation. The traditional multilingualism in indigenous languages in the present oldest generation has given way to bilingualism with the English-based creole Tok Pisin in an intermediate generation, then monolingualism in Tok Pisin, with perhaps English from schooling, in a third generation (46). With Greenberg’s language diversity index (the probability that an individual does not share the same language with another randomly selected individual) approaching 0.989 (Dataset S1), the languages of PNG are too localized to be practical for wider communication. Unfortunately, we have shown that ethnobiological knowledge is closely correlated with indigenous language skills and therefore equally at risk.

The factors predicting language and biocultural knowledge attrition in our models are determined by the factors considered desirable in contemporary PNG society, such as education, cash economy, ease of travel, and skills demanded for employment, or they are a consequence of economic development, such as urbanization, which also leads to mixed-language marriages. These are powerful forces making the preservation of traditional knowledge difficult. In 2013, PNG abandoned a decades-long experiment in allowing local communities to deliver early childhood education in local indigenous languages, moving to an English-only plan (47). Further, children often leave their home village to pursue education, which can cause attrition of their indigenous language skills (48). PNG’s extraordinary linguistic diversity and overwhelmingly rural population pose a challenge for state-delivered education but have played an important role in the retention of vast biocultural knowledge that exists outside the education system.

The survival of most indigenous languages and traditional knowledge will be determined by factors other than their practicality. On the positive note, PNG communities prize language as a marker of group identity (24). A majority, 88%, of the students fluent in an indigenous language expressed their intention to teach it to their children, but only 8% were motivated by practicality for communication, while the others valued language as an important part of their culture. It is possible that biocultural knowledge is less consciously prized than language skills and therefore even more in danger of disappearing than indigenous languages (41).

New Guinea’s share of global linguistic diversity is more than twice as high as its share of biological diversity (5, 17). The nation’s linguistic and biological diversity continues to be extensively studied (13, 18), with some sustained efforts at protection (47, 49), but both local and international programs to document and support ethnobiological diversity remain limited (3, 21). A better synergy between traditional biological knowledge and formal biology, such as grassroots paraecologist programs, could reinvigorate the interest of indigenous communities in their ethnobiological heritage as well as in the preservation of linguistic and biological diversity (40, 41, 17, 21).

## Material and Methods

### Language skills and ethnobiological knowledge variables

We surveyed students attending upper secondary school (grades 11 and 12) at 30 of the 123 secondary schools in PNG from 6^th^ April 2015 to 14^th^ November 2018. The schools were selected to represent both rural and urban locations in the lowland and highland regions from several provinces, comprising areas with both low and high language diversity (SI Appendix, Fig. S5, Table S2). The students completed tests of indigenous language skills and ethnobiological knowledge and a questionnaire on their family background, skills, and lifestyle (Dataset S3). All surveys were voluntary, anonymous and with informed consent given by all participants. They were conducted at schools and attained 100% participation, eliminating the problem of self-selection, when poor speakers may be reluctant to volunteer for language tests (Fig. 1B).

Indigenous language skills were quantified by two variables: [L1] the number of body parts, from a set of 24 that included both frequently and rarely used terms, named by students from photographs (50), and [L2] the student’s self-assessment on a four-point scale: 0 - no language skills; 1 - passive understanding; 2 - speaking but poorly; and 3 - fluent language use. Students speaking more than one indigenous language were assessed for the language they knew best. Language skills measures based on self-assessment (L2) can be biased (51). Individual respondents may have different ideas about what it means to be “fluent”. Younger people may consider themselves less linguistically fluent than elders because they have less cultural knowledge. However, these potential biases are unlikely to be important since the L1 and L2 variables are closely correlated in our study (SI Appendix, Fig. S6).

Ethnobiological knowledge was quantified by two variables: [E1] the number of bird species named in an indigenous language, from a set of images of 10 species, and [E2] the number of plant species freely listed with their indigenous names and traditional uses other than food (10 species maximum). For birds, the students completed two sets including, respectively, 10 lowland and 10 montane species, all geographically widespread. Each selection included a range of species, from widely known and easily recognizable ones (e.g., bird of paradise and cassowary species), to more difficult ones. The set with the higher score was used for each student, so as not to penalize students from any geographic location. We combined the image identification for birds with free listing for plants in order to obtain more comprehensive ethnobiological information as each method of data collection has its own strengths and biases (20, 52). The ethnobiological knowledge measures focused on indigenous species names for birds and plants, because knowledge of these names is a prerequisite for learning traditional information associated with individual species. Tok Pisin does not have detailed animal or plant taxonomies, and those available in English are not widely used in PNG. For plants, some students listed species by their Tok Pisin or English names only, when they did not know their indigenous names. These data were analyzed separately. Our tests, limited to 10 bird and 10 plant species, did not explore the full scope of ethnobiological knowledge which often includes several hundred species (20, 43). With their focus on students, they were also not designed to capture improvements of knowledge with age that often take place for people who are immersed in the relevant cultural and natural environment (53).

We used 21 independent variables (details in SI Appendix) to explain language skills and ethnobiological knowledge, categorized into four classes:

[A] Language traits: [A1] Language population size: We estimated the number of language users by interpolating or extrapolating the number listed in the *Ethnologue* database (1) to the year 2000 (Dataset S1). [A2-A3] Language status: We used either detailed EGIDS categories [A2] as given for each language in *Ethnologue* (1), or the language status [A3] classified as endangered (EGIDS 6b to 10) or not (EGIDS 1 – 6a). [A4] Geographic region: The location of the language in one of the four geographic and administrative regions of PNG (1 - Highlands, 2 - Momase, 3 - Southern and 4 – Islands) is used as a categorical variable to examine geographic differences in language skills. [A5] Elevation: Each language was characterized by its median elevation (in m asl, log transformed), obtained from the *Ethnologue* (1) language maps. [A6] Bible translation: Bible translations are typically the only written literature in indigenous languages that are used by their speakers.

[B] Family socio-economic traits: [B1] Urbanization: The student’s childhood place of residence is: 1 – village, 2 – government outpost, 3 – town or city. [B2] Remoteness: The student’s childhood place of residence can be accessed by: 1 – road, 2 – boat (no road), 3 – plane (no road or boat), 4 – only on foot. [B3] Parents’ education: The highest education reached by either of the parents: 1 – no school, 2 – lower primary (1^st^ – 6^th^) grade, 3 – higher primary (7^th^ – 8^th^) grade, 4 – lower secondary (9^th^ - 10^th^) grade, 5 – higher secondary (11^th^ – 12^th^) grade, 6 – any tertiary education. [B4] Parents’ employment: The highest employment category reached by either of the parents: 1 – subsistence farming, 2 – cash crop farming, 3 – salaried job or small business.

[C] Family language use: [C1] Parents’ language fluency: The L2 scores were assessed by the respondents for their parents; the higher of the mother’s and father’s scores was used. [C2] Home language use: 1 – indigenous language (alone or with other languages, including Tok Pisin and English), 2 – Tok Pisin only, 3 – English (alone or with Tok Pisin). [C3] Parents’ languages: 1 – mother and father speak the same indigenous language, 0 – the family is linguistically mixed.

[D] Student traits: [D1] Gender: 1 – female, 0 – male. [D2-D3] Grade 10 test results from English (D2) and Mathematics (D3). [D4-D6] Traditional and contemporary technical skills: Students self-assessed their skills (as 0 – none, 1 – poor, 2 – good) at five traditional tasks (D4): hunting, fishing, growing staple crops, building a house from forest materials, and using plants to treat fever, as well as at two contemporary technical tasks (D5): using a mobile phone and using a computer. The difference between traditional and contemporary technical skills was used as an additional explanatory variable (D6).

[D7] Best friend’s language: The participant’s best friend speaks the same (1) or a different (0) indigenous language as the informant. This variable was used as a proxy for the proportion of mixed language families likely to be formed by the surveyed students in the future (C3), viewed as a predictor of language skills for the generation of the students’ children. We compared the D7 values for each surveyed school with the expected proportion of best friends speaking the same language as the informant, with the assumption that students choose their friends at school and do so irrespective of the indigenous language they speak. [D8] Teaching indigenous language: The intention to teach one’s children an indigenous language (1 – yes, 0 – no), from those who have the language skills to do so, with a pre-defined list of five motivations to justify this choice: no, because (i) the indigenous language belongs to an old culture, or (ii) it is not a useful skill for my child; yes, because (i) everyone in my village/town does it, (ii) it is a useful skill for my children, or (iii) it is part of my culture. This variable was not used for GLM models.

### Data verification

Identification of the indigenous language used by each respondent was often difficult (details in SI Appendix). We were able to identify the indigenous language for 6,190 from 8,708 respondents. Both the complete and the verified data sets give similar results for the language skills (L1, L2) and ethnobiological knowledge (E1, E2); only verified data were used in the analysis. We also verified the body part test results, as detailed in the SI Appendix.

### Language skills and ethnobiological knowledge analysis

The data used for analysis are provided in Dataset S2. We used generalized linear mixed models to assess the effect of the four classes of variables on the language skills of the students. The response variable was the number of correct/incorrect body parts identified by students in their indigenous language (L1). The probability of getting correct responses was modelled as a binomial variable, with students and individual languages treated as random variables in all models. Except for A4, all other potential predictors are either binary or ordinal variables, allowing us to model these variables as numeric (and A1 as natively numeric), with orthogonal polynomials of order *N* – 1 representing the number of levels in each variable. This approach is equivalent to representing contrasts in a categorical variable but allows for numeric extrapolation of non-integer values.

We employed a hierarchical model selection approach using the Akaike Information Criterion (AIC) (54) to compare the fit among candidate models. First, we used model selection separately for each class (A – D) of predictor variables. For the variables that had more than one level (except categorical A4), we built models with polynomials of different order, from N – 1 levels to a simple linear relationship. For each class, we considered all the variable combinations within that class, using different polynomial orders when applicable. When more than one variable represented alternative expression of the same factor (A2 vs. A3, D6 vs. D4 and D5), we excluded models that included these variables together. In order to make interpretations easier, avoid inflation in the number of candidate models, and limit degrees of freedom in the models, we did not consider interactions among the variables. In the end, we obtained, for each class, one best-performing model with the optimal set of variables belonging to that class (SI Appendix, Table S2). Subsequently, we combined the variables from each of these class-specific, best-performing models to test whether different classes of variables acted jointly on language skills. We built these models by combining all the variables from the best-performing model in each class into models with two, three, and all four classes, in all possible combinations, again with no interactions (SI Appendix, Table S3).

Once we obtained the best performing, overall model, we investigated the relative role of each class and individual variable by calculating how much the AIC value was increased by removing the focal variable or class from the full model. In case any variable came out with a non-significant marginal effect (using a threshold of 2 points of AIC) at this stage, we removed it from the final model, as such loss of effect would be due to a higher predictive power in other correlated variables from another class. In addition, we assessed the direction and shape of the effects for individual predictor variables (Fig. 3). We predicted the response variable while varying each of the predictor variables in the best performing model across its range, while keeping the other predictors at their original mean values across the whole population of test scores. This procedure was not possible for geographic regions (A4), which is categorical. We kept the A4 values fixed on the most abundant region when predicting the effect of the other variables, because an average does not apply to this categorical variable.

We also used the generalized linear mixed models to analyze the ethnobiological knowledge of students (E1 and E2). We used the same variables and model building strategy as for L1, except that we omitted the Language traits (A) class of variables and added language skills (L1) as a new independent variable.

### Language skills extrapolation

The empirical relationships among the predictor variables included in the best model were used to extrapolate language skills (L1) for hypothetical populations characterized by values for the model parameters different from the observed: for the 18-20-year-olds in PNG at present (Model L1A), characterized by parameters extrapolated 30 years into the future (Model L1B), and assuming PNG reaches the current mean socio-economic parameters for the Lower-Middle Income Countries (55) (Model L1C). We used the estimated parameters for the effects of each variable from the best model described above to predict the response variable for different values of the predictor variables in these populations. Because variable A4 is categorical, we made separate predictions for each geographic region, and then made an average prediction by weighting the predicted value for each region by its proportion in the overall population.

In Model L1A, we used parameter values characterizing 18-20-year-olds in PNG for urbanization (B1), parents’ education (B3), parents’ employment (B4), geographic region (A4), and language status (A3) (1, 29, 34, 37) (SI Appendix, Table S4). These variables were also used to adjust the remaining variables for which PNG-wide data were not available. For instance, there are no country-wide data for remoteness (B2), but the distribution of remoteness values differs between two levels of urbanization – village and town/city. The remoteness variable was therefore adjusted as a weighted mean between village and town/city values, reflecting the change in the share of village residents from 58% in the original data to 87% in Model L1A. More complex adjustments included several explanatory variables, using distributions of the adjusted variable for all possible combinations of their values. In Model L1A, in addition to remoteness adjusted by urbanization, we adjusted four variables (C2, C3, D4, and D5) by a combination of urbanization and parental education (SI Appendix, Table S4). The parameters for the L1B and L1C models, extracted from literature (1, 34, 36, 37, 56) and adjusted using other variables, appear in SI Appendix, Table S4. Finally, we adjusted the proportion of respondents fluent in an indigenous language (L2 = 3) from the student respondents in our study to the entire 18-20-year-old cohort in PNG and used this approach to estimate the proportion of fluent speakers in the next generation, both for students (Model L2B) and entire 18-20-year-old cohort in PNG (Model L2C) (SI Appendix, Table S4).

### Data availability

All data used for the analysis are included in the article, the SI Appendix and Datasets S1-S2.

## ACKNOWLEDGEMENTS

We thank students, teachers, and heads of the 30 secondary schools that participated in the study; the PNG Department of Education for support; the PNG National Research Institute, particularly G. Kaipu, for permitting the research; and the New Guinea Binatang Research Center for logistical support. G. Luke, M. Sisol, M. Kigl and E. Messimato provided information on individual languages; K. Kama assisted data collection, J. Leps advised on statistical analysis. C. Volker commented on the manuscript. The New Guinea Binatang Research Center has provided the ethics oversight of the study, including data collection. The study was supported by the Czech Science Foundation (18-23889S), The European Research Council (669609), The Christensen Fund (2017-10237), and The Darwin Initiative for the Survival of the Species (DIR25S1\100123).

## References

1. D. M. Eberhard, G. F., Simons, C. D. Fennig, Eds., Ethnologue: Languages of the World (SIL International, ed. 23, 2020).

2. W. J. Sutherland, Parallel extinction risk and global distribution of languages and species. Nature 423, 276–279 (2003).

3. R. Cámara–Leret, Z. Dennehy, Indigenous knowledge of New Guinea’s useful plants: a review, Econ. Bot. 73, 405–415 (2019).

4. L. Maffi, Linguistic, cultural, and biological diversity. Annu. Rev. Anthropol. 29, 599–617 (2005).

5. V. Novotny, P. Drozd, Size distribution of conspecific populations: peoples of New Guinea. Proc. R. Soc. Lond., Biol. Sci. 267, 947–952 (2000).

6. L. Campbell, E. Okura, “New knowledge produced by the Catalogue of Endangered Languages” in Cataloguing the World’s Endangered Languages, L. Campbell, A. Belew, Eds. (Routledge, 2018), pp. 79–84.

7. M. L. Landweer, Methods of language endangerment research: a perspective from Melanesia. Int. J. Sociol. Lang. 214, 153–178 (2012).

8. T. Amano, B. Sandel, H. Eager, E. Bulteau, J-C. Svenning, B. Dalsgaard, C. Rahbek, R. G. Davies, W. J. Sutherland, Global distribution and drivers of language extinction risk. Proc. R. Soc. B, 281, 20141574 (2014).

9. K. Prochazka, G. Vogl, Quantifying the driving factors for language shift in a bilingual region. PNAS 114, 4365–4369 (2017).

10. L. Bromham, X. Hua, C. Algy, F. Meakins, Language endangerment: a multidimensional analysis of risk factors. J. Lang. Evol., 5, 75–91 (2020).

11. S. Aswani, A. Lemahieu, W. H. H. Sauer, Global trends of local ecological knowledge and future implications. PLoS ONE 13, e0195440 (2018).

12. T. Furusava, Changing ethnobotanical knowledge of the Roviana people, Solomon Islands: Quantitative approaches to its correlation with modernization. Hum. Ecol. 37, 147–159 (2009).

13. B. Palmer, The Languages and Linguistics of the New Guinea Area: A Comprehensive Guide. (De Gruyter Mouton, 2017).

14. H. Hammarström, R. Forkel, M. Haspelmath, S. Bank, Sebastian, Glottolog 4.2.1. (Max Planck Inst. Science of Human History, Jena (March 2020); http://glottolog.org.

15. J. Vaughan, R. Singer, Indigenous multilingualisms past and present. Lang. Commun. 62, 83–90 (2018).

16. S. A. Wurm, “Threatened languages in the Western Pacific Area from Taiwan to, and including, Papua New Guinea” in Language Diversity Endangered, M. Brenzinger, Ed. (Mouton de Gruyter, 2007), pp. 374–390.

17. V. Novotny, K. Molem, An inventory of plants for the land of the unexpected. Nature 584, 531–533 (2020).

18. V. Novotny, P. Toko, “Ecological research in Papua New Guinean rainforests: insects, plants and people”, in J. E. Bryan, P. L. Shearman, Eds., The State of the Forests of Papua New Guinea 2015. Measuring change over period 2002-2014 (University of Papua New Guinea, 2015), pp. 71–85.

19. P. Mühlhäusler, “The development of the life form lexicon in Tok Pisin” in Degrees of Restructuring in Creole Languages, I. Neumann-Holzschuh, E. W. Schneider, Eds. (J. Benjamins, 2000), pp. 337–360.

20. B. Berlin, Ethnobiological classification. Principles of Classification of Plants and Animals in Traditional Societies (Princeton Univ., 1992).

21. L. Maffi, E. Woodley, Biocultural Diversity Conservation. A Global Sourcebook (Earthscan, 2010).

22. S. Zent, “Processual perspectives on traditional environmental knowledge. Continuity, erosion, transformation, innovation”, in R. Ellen, S. J. Lycett, S. E. Johns, Eds., Understanding Cultural Transmission in Anthropology: A Critical Synthesis (Berghahn Books, 2013), pp. 213–265.

23. D. A. Kulick, Death in the Rainforest. How a Language and a Way of Life Came to an End on Papua New Guinea (Algonquin Books, 2019).

24. K. M. Sumbuk, “Papua New Guinea’s languages: will they survive?” in Language Diversity in the Pacific: Endangerment and Survival D. D. Cunningham, P. D. E. Ingram, P. K. Sumbuk, Eds. (Multilingual Matters, 2006), pp. 85–96.

25. A. Y. Aikhenvald, Language contact along the Sepik River, Anthropol. Linguist. 50, 1–66 (2008).

26. F. Lüpke, Uncovering small-scale multilingualism. Crit. Multilingualism Stud. 4, 35–74 (2016).

27. A. Y. Aikhenvald, “Language endangerment in the Sepik area of Papua New Guinea” in Lectures on Endangered Languages: 5. Endangered Languages of the Pacific Rim, O. Sakiyama, F. Endo, Eds. (ELPR, 2004), pp. 97–142.

28. S. Romaine, Language, Education, and Development: Urban and Rural Tok Pisin in Papua New Guinea, (Clarendon Press, 1992).

29. Worldometers, Papua New Guinea Population (March 2020); https://www.worldometers.info/world-population/papua-new-guinea-population/.

30. G. Koczberski, G. N. Curry, B. Imbun, Property rights for social inclusion: Migrant strategies for securing land and livelihoods in Papua New Guinea. Asia Pacif. Viewp. 50, 29–42 (2009).

31. M. P. Lewis, G. F. Simons, Assessing endangerment: expanding Fishman’s GIDS. Rev. roum. linguist. 55, 103–120 (2010).

32. R. van den Berg, Scripture Use Research and Ministry, (SIL International, 2020).

33. P. K. Austin, J. Sallabank, “Introduction” in Cambridge Handbook of Endangered Languages, P. K. Austin, J. Sallabank, Eds., (Cambridge Univ., 2011), pp. 3–24.

34. National Statistical Office (NSO), Papua New Guinea Demographic and Health Survey 2016-18 (NSO, 2019).

35. S. E., Vollset, E. Goren, C-W. Yuan, J. Cao, A. E. Smith, T. Hsiao, C. Bisignano, G. S. Azhar, E. Castro, J. Chalek, A. J. Dolgert, T. Frank, K. Fukutaki, S. I. Hay, R. Lozano, A. H. Mokdad, V. Nandakumar, M. Pierce, M. Pletcher, T. Robalik, K. M. Steuben, H. Y Wunrow, B. S. Zlavog, C. J. L. Murray, Fertility, mortality, migration, and population scenarios for 195 countries and territories from 2017 to 2100: a forecasting analysis for the Global Burden of Disease Study. Lancet 396, 1285–1306 (2020).

36. United Nations, World Urbanization Prospects: The 2018 Revision, (United Nations, 2019).

37. National Statistical Office (NSO). 2009-2010 Papua New Guinea Household Income and Expenditure Survey, (NSO, 2011).

38. A. Si, Patterns in the transmission of traditional ecological knowledge: a case study from Arnhem Land, Australia. J. Ethnobiol. Ethnomed. 16, 52 (2020).

39. E. S. Hunn, A Zapotec Natural History: Trees, Herbs, and Flowers, Birds, Beasts, and Bugs in the Life of San Juan Gbëë (Arizona Univ. Press, 2008).

40. V. Novotny, G. D. Weiblen, S. E. Miller, Y. Basset, The role of paraecologists in 21st century tropical forest research, in M. D. Lowman, T. D. Schowalter, J. F. Franklin, Eds., Methods in Forest Canopy Research (Univ. of California, 2012), pp. 154–157.

41. I. S. Majnep, R. Bulmer, Birds of my Kalam Country. (Auckland Univ. & Oxford Univ., 1977)

42. R. Bulmer, Why is the cassowary not a bird? A problem of zoological taxonomy among the Karam of the New Guinea Highlands. Man NS 2, 5–25 (1967).

43. R. O. Gardner, Plant names of the Kalam (Upper Kaironk Valley, Schrader Range, Papua New Guinea). Rec. Auckland Mus. 47, 5–50 (2010).

44. O. Potterat, M. Hamburger, Morinda citrifolia (Noni) fruit - phytochemistry, pharmacology, safety. Planta Med. 73, 191–199 (2007).

45. P. K. Austin, J. Sallabank, Eds. The Cambridge Handbook of Endangered Languages (Cambridge Univ., 2011).

46. A. Y. Aikhenvald, “Traditional multilingualism and language endangerment” in Language Endangerment and Language Maintenance, D. Bradley, M. Bradley, Eds. (Routledge, 2002), pp. 24–33.

47. P. Paraide, Challenges with the implementation of vernacular and bilingual education in Papua New Guinea. Contemp. PNG Stud.: DWU Res. J. 21, 44–57 (2014).

48. B. Köpke, Neurolinguistic aspects of attrition. J. Neuroling. 17, 3–30 (2004).

49. V. M. Adams, N. Dimitrova, H. P. Possingham, J. R. Allan, C. D. Kuempel, N. Peterson, A. Kaiye, M. Keako, V. J. D. Tulloch, Scheduling incremental actions to build a comprehensive national protected area network for Papua New Guinea. Cons. Sci. Pract. 3, e354 (2021).

50. W. O. O’Grady, A. J. Schafer, J. Perla, O.-S. Lee, J. Wieting, A psycholinguistic tool for the assessment of language loss: the HALA Project. Lang. Doc. Conserv. 3, 100–112 (2009).

51. C. Grinevald, M. Bert, “Speakers and communities” in Cambridge Handbook of Endangered Languages, P. K. Austin, J. Sallabank, Eds. (Cambridge Univ., 2011), pp. 45–65.

52. V. Reyes-García, N. Martí, T. McDade, S. Tanner, V. Vadez, Concepts and methods in studies measuring individual ethnobotanical knowledge. J. Ethnobiol. 27, 182–203 (2007)

53. J. Koster, R. McElreath, K. Hill, D. Yu, G. Shepard Jr., N. van Vliet, M. Gurven, B. Trumble, R. B. Bird, D. Bird, B. Codding, L. Coad, L. Pacheco-Cobos, B. Winterhalder, K. Lupo, D. Schmitt, P. Sillitoe, M. Franzen, M. Alvard, V. Venkataraman, T. Kraft, K. Endicott, S. Beckerman, S. A. Marks, T. Headland, M. Pangau-Adam, A. Siren, K. Kramer, R. Greaves, V. Reyes-García, M. Guèze, R. Duda, Á. Fernández-Llamazares, S. Gallois, L. Napitupulu, R. Ellen, J. Ziker, M. R. Nielsen, E. Ready, C. Healey, C. Ross, The life history of human foraging: Cross-cultural and individual variation. Sci. Adv. 6, eaax9070 (2020).

54. K. P. Burnham, D. R. Anderson, Model Selection and Multimodel Inference. A Practical Information-Theoretic Approach (Springer, 2nd Ed., 2002).

55. World Bank, Country Classification (March 2020); https://datahelpdesk.worldbank.org/knowledgebase/topics/19280-country-classification.

56. World Bank, Data Catalog: Educational Statistics (March 2020); https://datacatalog.worldbank.org/dataset/education-statistics.

