## Supplemental Material for "The world’s hotspot of linguistic and biocultural diversity under threat"

### Supplementary Methods and Materials

#### Language skills and ethnobiological knowledge variables

Methodological details for selected variables:

[A] Language traits: [A1] Language population size: The number of language users was estimated by adjusting the number listed in the *Ethnologue* database (1), interpolated or extrapolated to the year 2000 using annual population growth rates for PNG (2). This standardization was necessary, because the *Ethnologue* estimates date from 1971 – 2019 for individual languages, most often using the PNG National Census data from 2000 (3), and the Summer Institute of Linguistics estimates date from 2003 (Dataset S1). [A2] Language status: We used detailed EGIDS categories as given for each language in *Ethnologue* (1): 1 – EGIDS 3, 2 – EGIDS 4, 3 – EGIDS 5, 4 - EGIDS 6a, 5 – EGIDS 6b, 6 – EGIDS 7 to 10. [A5] Elevation: Each language was characterized by its median elevation (in m, log transformed), obtained from the *Ethnologue* (1) language maps overlaid on geographic maps using Zonal Statistics tool ArcGIS Pro on the Shuttle Radar Topography Mission (SRTM) elevation dataset (4) with spatial resolution 3" latitude x 3" longitude. [A6] Bible translation: We used the lists of languages with at least partial Bible translations (1, 5), but we do not have information on the use of these translations by the surveyed students.

[D] Student traits: [D2-D3] Grade 10 test results from English (D2) and Mathematics (D3): 1 – distinction, 2 – credit, 3 – upper pass, 4 – pass or fail. [D6] The scores were summed within traditional (maximum 10 points) and contemporary technical (maximum 4 points) activities, the totals were rescaled to a 0 – 1 range, and the difference between traditional and contemporary technical skills was used as an explanatory variable. [D7] Best friend's language: We compared the observed values for each surveyed school with the expected proportion of best friends speaking the same language as the informant, calculated as $\sum_{i=1}^{n} p_i^2$, where $p_i$ is the proportion of students speaking language $i$ in the surveyed school. This probability assumes that students choose their friends at school and do so irrespective of the indigenous language they speak. The overall random expectation of the best friend's language was calculated as the average for all 30 schools surveyed, weighted by their student numbers.

#### Data verification

Identification of the indigenous language used by each respondent was often difficult. The respondents were often unaware of the name used for their indigenous language by linguists (1) and gave alternative, local names for languages as well as for individual dialects. Further, geographic distribution of many languages in PNG is poorly known, so that language maps remain approximate (1). Villages in PNG also often change their location or name. We therefore integrated information on the language name given by each respondent, the respondent's birthplace, the language and birthplace of the respondent's parents, and the results of the language test naming individual body parts (L1), in order to identify the indigenous language used by the respondent.

We verified the body part test results from 1,990 respondents (32% of the total) speaking the Melpa, Kuman, Enga, and Amele languages (i.e., four of the six languages represented by >100 respondents) with the help of native speakers. The assessment of the responses in other languages was made difficult by the lack of vocabulary lists for many languages (6) and by dialectical differences, which are often poorly documented. Most respondents do not write in their indigenous language, resulting in widely variable spelling in their written responses. The terms for some body parts included in the test may not be widely used in some languages, resulting in particularly high error rates for them. The 47,760 test questions (24 body parts for 1,990 respondents) yielded an answer rate of 81%, including 63% correct answers, 16% answers that referred either to a related body part, or a wider anatomical area (e.g., hand instead of wrist, or toe instead of toenail), and 2% of the answers that were entirely wrong. These data from a few common languages were used to develop a universal protocol applied to all languages. We considered the correct and partly incorrect or vague responses as valid since they reflected some knowledge of the language, as opposed to the entirely incorrect and non-responses. This strategy was also used for the remaining 388 languages, as far as possible. When we were unsure about the correct term, we accepted the response given by a majority of respondents. In languages represented by a single or a few respondents and lacking linguistic information, we accepted the responses provided as valid, since our detailed analysis of the four common languages indicated that this was predominately the case. We used the same approach to verify indigenous bird names in the tests. The indigenous names of plant species that were freely listed were all accepted as correct, since it was impossible to verify them across 392 languages in our data, most of which are ethnobotanically undocumented (7). Our methods of data verification likely somewhat overestimated language skills and ethnobiological knowledge of the respondents by accepting some erroneous responses as valid. On the other hand, many respondents could be unaccustomed to write in their indigenous language, which could have negatively affected their written test results.

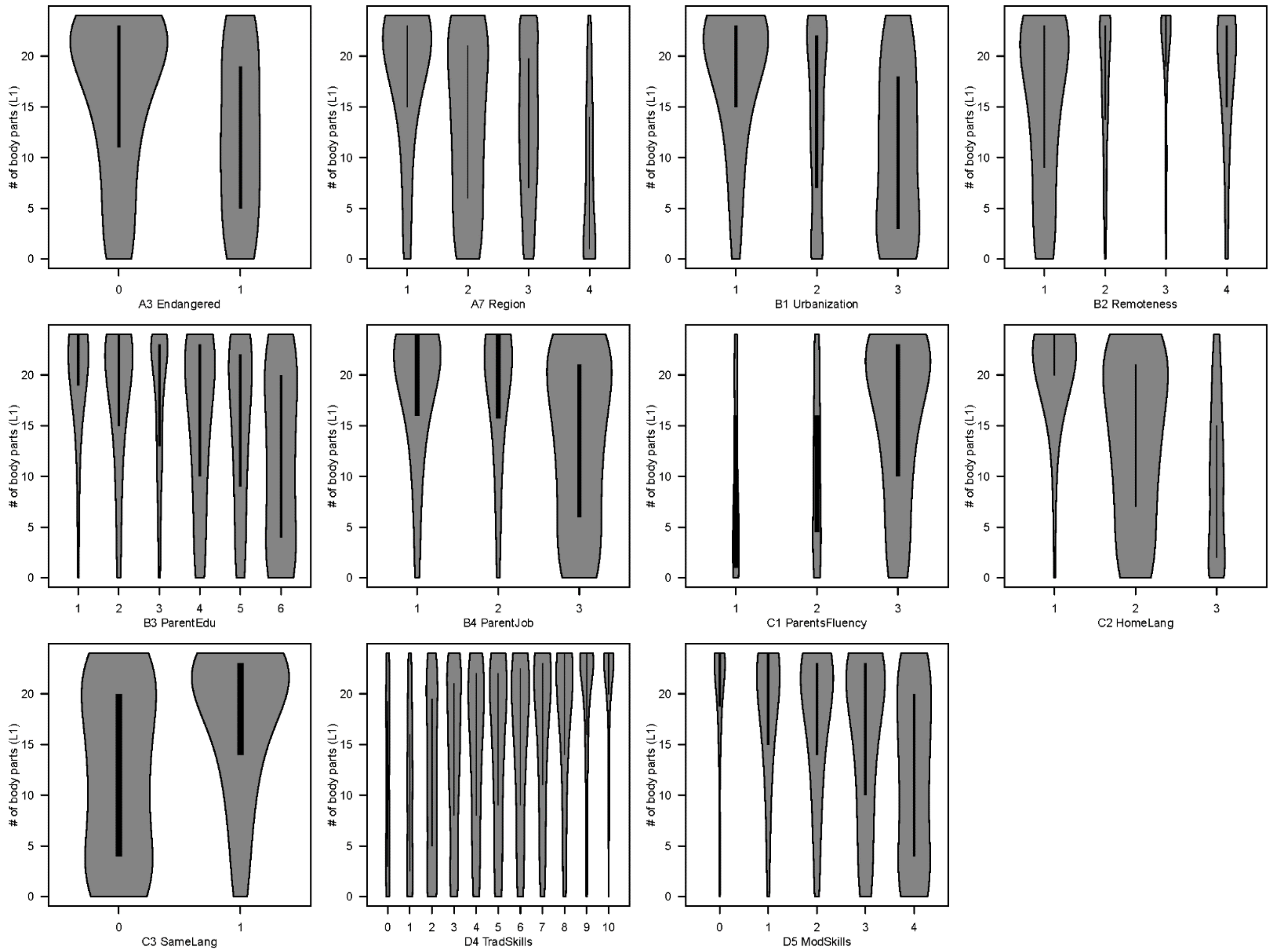


**Fig. S1.** Effect of the independent variables explaining the indigenous language skills (L1) of students from PNG. Each bar shows the density distribution of the L1 response variable for a given level of the predictor variable, with the width of the bar proportional to the number of students belonging to that class. See Materials and Methods for details on the variables.

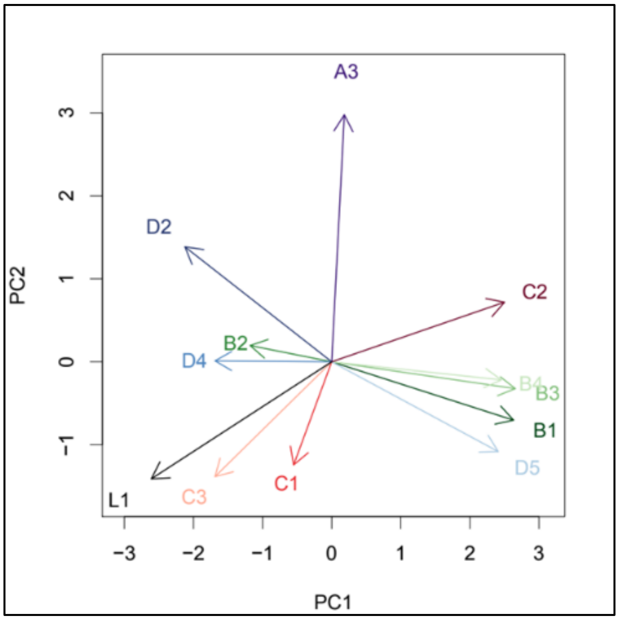


**Fig. S2.** Correlation of independent variables used to explain language skills. PCA ordination of the 12 independent variables included in the model explaining language skills (L1). All variables were considered numerical, and only linear correlations are visible in this PCA. Variable A4 was excluded as categorical. Arrows represent the direction in which each variable increases along the two PCA axes; their color coding is the same as in Fig. 3. PC1 explained 28.1% and PC2 explained 10.3% of the total variation in the data. See Materials and Methods for the description of variables.

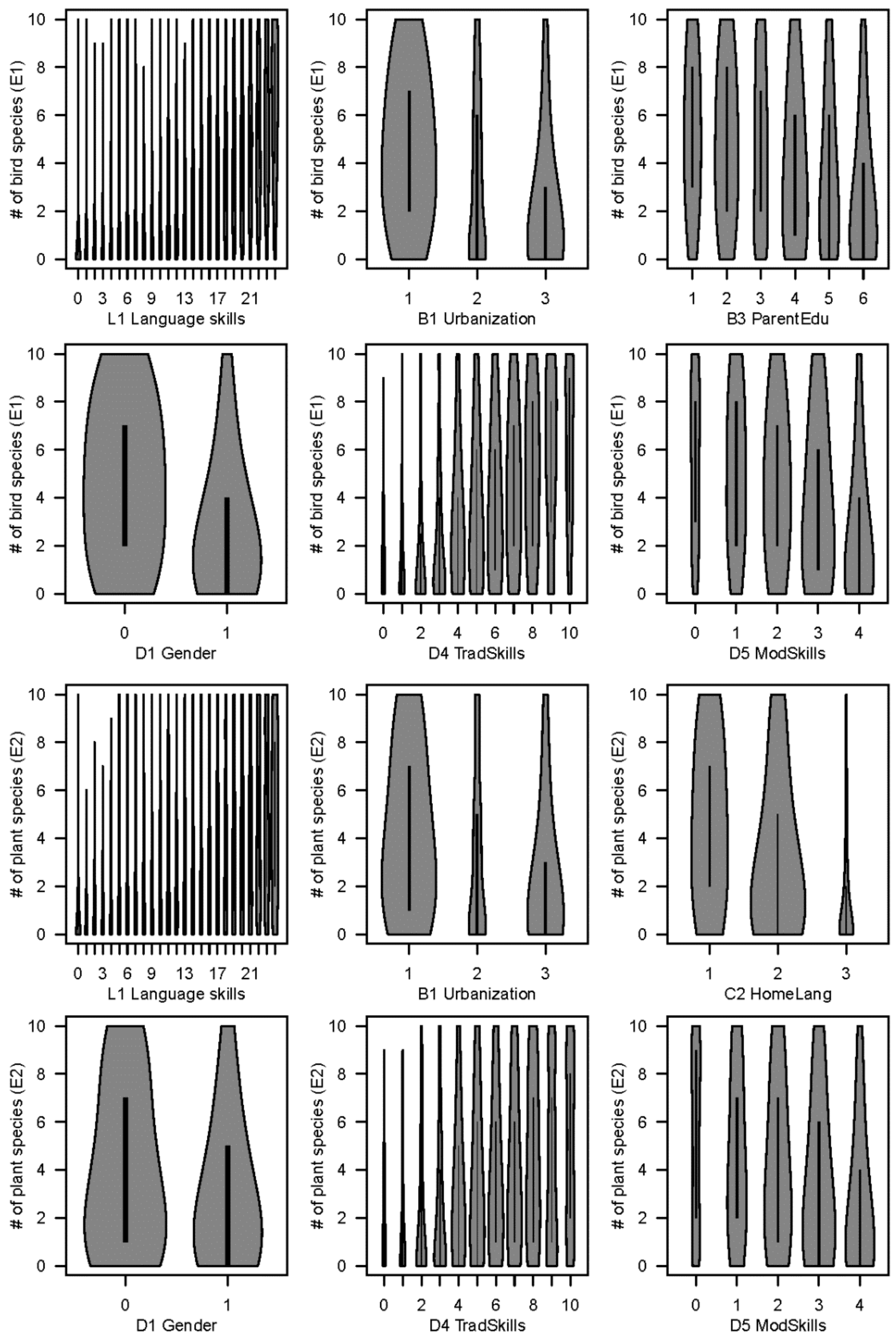


**Fig. S3.** Effect of the independent variables explaining the ethnobiological knowledge (E1, E2) of students from PNG. Each bar shows the density distribution of the E1 or E2 response variable for a given level of the predictor variable, with the width of the bar proportional to the number of students belonging to that class. See Materials and Methods for details on the variables.

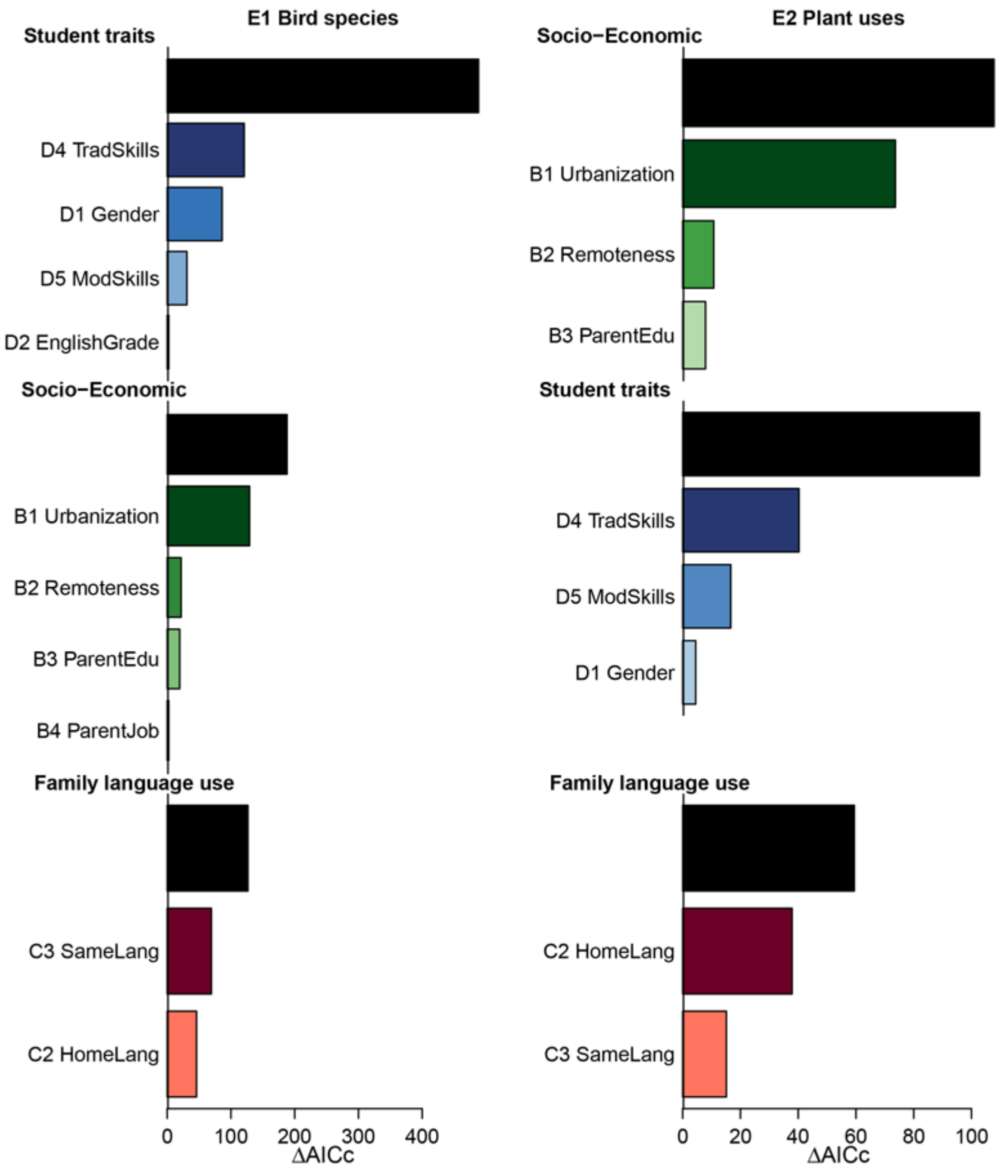


**Fig. S4.** Effects of language and socio-economic factors on ethnobiological knowledge. Generalized linear mixed models (GLMM) describe variability in the knowledge of bird species (E1, left) and traditional plant uses (E2, right) of students. The models incorporated 11 fixed variables divided into three classes: (B) Socio-economic traits: student's birthplace urbanization (B1) and remoteness (B2), parents' education (B3) and parent's employment (B4); (C) Family language use: parents' language skills (C1), home language use (C2), and whether parents speak the same first language (C3); and (D) Student traits: gender (D1), English skills (D2), traditional skills (hunting, fishing, growing food, house building, medicinal plants) (D4), contemporary technical skills (mobile phone and computer use) (D5). The variables were selected within each class before being included in a global model. The bars show the AIC improvement due to addition of each group (black) and each variable within each group into a model that include all other variables, quantifying the marginal effect of each class/variable. Details on the variables are in Materials and Methods. Only significant variables are shown.

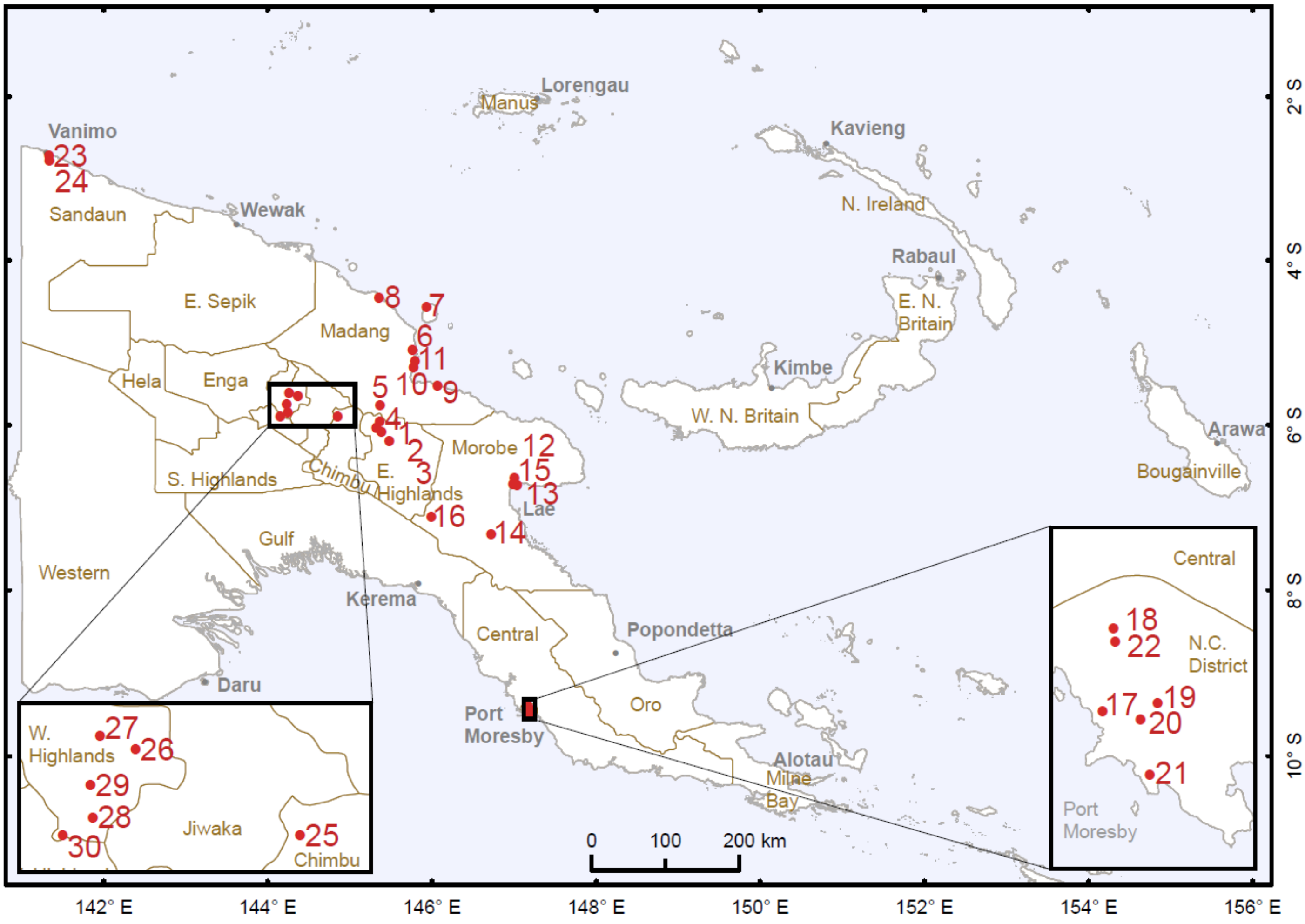


**Fig. S5.** Location of the 30 secondary schools surveyed in the study. See Table S2 for the details on individual schools.

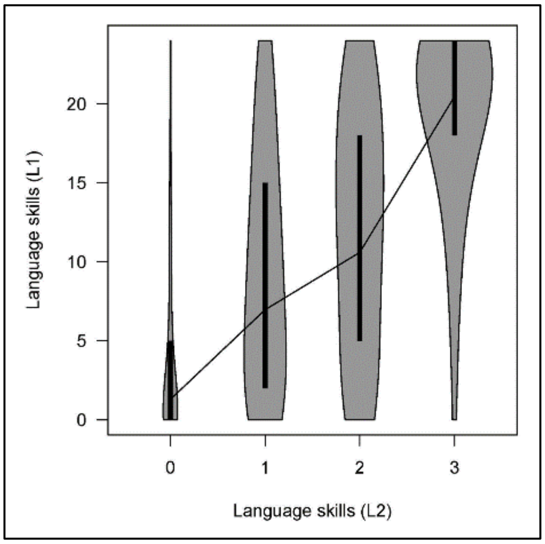


**Fig. S6.** Correlation between two language skills measures: self-assessed language fluency (L2) and the number of body parts named (L1) for the 6,190 students (Spearman $r = 0.58$, $P < 0.001$).

**Table S1.** Socioeconomic and language parameters for respondents. See Materials and Methods for details of variables A1 - D8. Based on the survey of N = 6,190 secondary school students.

| Var# | Variable | Value |
|---|---|---|
| A1 | Language population size [median, Q1-Q3] | 3093 [1350, 8500] |
| A3 | Students speaking endangered languages, % | 13.9 |
| A6 | Students speaking languages with a Bible translation, % | 84.0 |
| B1 | Students who spent childhood in a village, % | 64.7 |
| B2 | Student's childhood residence accessible by road % | 80.4 |
| B3 | Families having ≥1 parent with secondary of higher education, % | 63.2 |
| B4 | Families with subsistence agriculture income only, % | 30.9 |
| C2 | Families using indigenous language at home, % | 30.0 |
| C3 | Families with parents speaking the same indigenous language, % | 63.1 |
| D1 | Female students, % | 41.0 |
| D4 | Students with good hunting skills, % | 22.4 |
| D4 | Students with good fishing skills, % | 39.3 |
| D4 | Students with good farming skills, % | 67.8 |
| D4 | Students with good house building skills, % | 28.1 |
| D4 | Students with good plant medicinal use skills, % | 31.0 |
| D5 | Students with good mobile phone use skills, % | 67.9 |
| D5 | Students with good computer use skills, % | 28.3 |
| D7 | Students speaking the same indigenous language as their best friend | 34.8 |
| D8 | Students able and wishing to teach indigenous language to their child | 87.7 |
| | Student's age [median, Q1-Q3) | 19 [18, 20] |
| | Student's no. of siblings [median, Q1-Q3] | 4 [3, 6] |
| | Families owning land, % | 91.7 |
| | Families owning cash crop plantation, % | 52.0 |
| | Families owning forest, % | 68.5 |
| | Families with access to electricity, % | 53.8 |

**Table S2.** Secondary schools surveyed in the study. The number of surveyed students, the number of indigenous languages spoken, and the percentage of students speaking the most common language are listed for each school, numbered as in Fig. S5 showing its geographic location.

| # | School | Province | Students | Languages | 1st lang % |
|---|---|---|---|---|---|
| 1 | Asaroka Lutheran Secondary | Eastern Highlands | 149 | 27 | 19 |
| 2 | Benabena Secondary | Eastern Highlands | 223 | 34 | 39 |
| 3 | Goroka Secondary School | Eastern Highlands | 44 | 20 | 23 |
| 4 | Kabiufa Secondary | Eastern Highlands | 137 | 27 | 18 |
| 5 | Braham Secondary | Madang | 104 | 48 | 13 |
| 6 | Good Shepherd Lutheran Secondary | Madang | 126 | 51 | 15 |
| 7 | Karkar Secondary | Madang | 228 | 48 | 28 |
| 8 | Malala Catholic Upper Secondary | Madang | 131 | 46 | 13 |
| 9 | Raikos Lutheran Secondary | Madang | 100 | 33 | 19 |
| 10 | Transgogol High | Madang | 37 | 18 | 35 |
| 11 | Tusbab Secondary | Madang | 205 | 72 | 12 |
| 12 | Bumayong Secondary | Morobe | 276 | 84 | 7 |
| 13 | Busu Secondary School | Morobe | 227 | 90 | 8 |
| 14 | Grace Memorial Secondary | Morobe | 212 | 62 | 17 |
| 15 | Lae National High | Morobe | 383 | 124 | 7 |
| 16 | Menyamya Secondary | Morobe | 182 | 34 | 25 |
| 17 | Badihagwa Secondary Technical | Nat. Capital Distr. | 43 | 28 | 9 |
| 18 | Gerehu Secondary | Nat. Capital Distr. | 157 | 52 | 11 |
| 19 | Gordons Secondary | Nat. Capital Distr. | 157 | 62 | 7 |
| 20 | Jubilee Catholic Secondary | Nat. Capital Distr. | 26 | 20 | 8 |
| 21 | Kila Kila Secondary | Nat. Capital Distr. | 74 | 38 | 12 |
| 22 | Port Moresby National High | Nat. Capital Distr. | 69 | 33 | 10 |
| 23 | Don Bosco Secondary | Sandaun | 117 | 45 | 24 |
| 24 | Vanimo Secondary | Sandaun | 173 | 60 | 21 |
| 25 | Kerowagi Secondary | Simbu | 495 | 27 | 76 |
| 26 | Kitip Secondary | Western Highlands | 346 | 25 | 82 |
| 27 | Kwip Dau Secondary | Western Highlands | 301 | 23 | 79 |
| 28 | Mount Hagen Secondary | Western Highlands | 483 | 36 | 66 |
| 29 | Paglum Adventist Secondary | Western Highlands | 189 | 35 | 50 |
| 30 | Togoba Secondary | Western Highlands | 422 | 17 | 63 |

**Table S3.** Model selection results for the candidate models assessing the role of language-related variables A1 – A6, socio-economic variables B1 – B4, family language use variables C1 – C3, and student-related variables D1 – D6 on the language fluency of students. dAICc = delta corrected Akaike Information Criterion, df = degrees of freedom. All models include Student and Language as random factors on the intercept. Note that D7 and D8 were not used in model building.

| Variable class | Models – Student trait variables | dAICc | df |
|---|---|---|---|
| Language traits | A1_LogN.2000 + A3_Endangered + A7_RegCode + A8_LogElevMedian | 0 | 9 |
| Language traits | A1_LogN.2000 + A3_Endangered + A6_PNGbible + A7_RegCode + A8_LogElevMedian | 0.72 | 10 |
| Language traits | A1_LogN.2000. + A2_Status + A6_PNGbible + A7_RegCode + A8_LogElevMedian | 3.08 | 13 |
| Language traits | A3_Endangered + A7_RegCode + A8_LogElevMedian | 3.27 | 8 |
| Language traits | A3_Endangered + A6_PNGbible + A7_RegCode + A8_LogElevMedian | 5.18 | 9 |
| Language traits | A1_LogN.2000 + A3_Endangered + A7_RegCode | 6.45 | 8 |
| Language traits | A3_Endangered + A7_RegCode | 6.91 | 7 |
| Language traits | A6_PNGbible + A7_RegCode + A8_LogElevMedian | 20.95 | 8 |
| Language traits | A1_LogN.2000 + A3_Endangered + A8_LogElevMedian | 35.78 | 6 |
| Language traits | Null (random factors + intercept) | 72.68 | 3 |
| Socio-economic | B1_Urbanization + B2_Isolation^2 + B3_ParentEdu + B4_ParentJob^2 | 0 | 9 |
| Socio-economic | B1_Urbanization + B2_Isolation^2 + B3_ParentEdu^2 + B4_ParentJob^2 | 0.96 | 10 |
| Socio-economic | B1_Urbanization + B2_Isolation^3 + B3_ParentEdu + B4_ParentJob^2 | 1.82 | 10 |
| Socio-economic | B1_Urbanization^2 + B2_Isolation^3 + B3_ParentEdu + B4_ParentJob^2 | 4.80 | 11 |
| Socio-economic | B1_Urbanization) + B2_Isolation^2 + B3_ParentEdu^2 + B4_ParentJob | 12.07 | 9 |
| Socio-economic | B1_Urbanization + B2_Isolation + B3_ParentEdu + B4_ParentJob^2 | 24.23 | 8 |
| Socio-economic | B1_Urbanization + B2_Isolation^2 + B3_ParentEdu | 37.19 | 7 |
| Socio-economic | B1_Urbanization + B3_ParentEdu + B4_ParentJob^2 | 64.94 | 7 |
| Socio-economic | B1_Urbanization + B2_Isolation^2 + B4_ParentJob^2 | 93.49 | 8 |
| Socio-economic | B2_Isolation^2 + B3_ParentEdu + B4_ParentJob^2 | 378.61 | 8 |
| Socio-economic | Null (random factors + intercept) | 992.17 | 3 |
| Language use | C2_HomeLang^2 + C1_ParentMaxFluency + C3_SameLang | 0 | 7 |
| Language use | C2_HomeLang^2 + C1_ParentMaxFluency^2 + C3_SameLang | 1.08 | 8 |
| Language use | C2_HomeLang + C1_ParentMaxFluency + C3_SameLang | 32.04 | 6 |
| Language use | C2_HomeLang + C3_SameLang) | 45.72 | 5 |
| Language use | C2_HomeLang^2 + C1_ParentMaxFluency | 249.60 | 6 |
| Language use | C3_SameLang | 533.92 | 4 |
| Language use | Null (random factors + intercept) | 917.26 | 3 |
| Student traits | D2_GradeEng^2 + D4_TradSkills + D5_ModSkills | 0 | 7 |
| Student traits | D2_GradeEng^3 + D4_TradSkills + D5_ModSkills | 0.94 | 8 |
| Student traits | D1_Gender + D2_GradeEng^3 + D4_TradSkills + D5_ModSkills | 2.08 | 9 |
| Student traits | D2_GradeEng^3 + D3_GradeMath^3 + D4_TradSkills + D5_ModSkills | 4.01 | 11 |
| Student traits | D1_Gender + D2_GradeEng^3 + D3_GradeMath^3 + D4_TradSkills + D5_ModSkills | 4.84 | 12 |
| Student traits | D2_GradeEng^3 + D6_Trad.Mod | 12.58 | 7 |
| Student traits | D2_GradeEng + D4_TradSkills + D5_ModSkills | 13.01 | 6 |
| Student traits | D1_Gender + D2_GradeEng^3 + D3_GradeMath^3 + D6_Trad.Mod | 17.00 | 11 |
| Student traits | D4_TradSkills + D5_ModSkills | 48.06 | 5 |
| Student traits | D1_Gender + D4_TradSkills + D5_ModSkills | 49.46 | 6 |
| Student traits | D1_Gender + D2_GradeEng^3 + D3_GradeMath^3 + D4_TradSkills | 199.85 | 11 |
| Student traits | D1_Gender + D2_GradeEng^3 + D3_GradeMath^3 + D5_ModSkills | 246.61 | 11 |
| Student traits | Null (random factors + intercept) | 750.43 | 3 |

**Table S4.** Model selection results for the candidate models assessing the combined role of the four variable classes A – D included in this study on the language fluency of students. dAICc = delta corrected Akaike Information Criterion, df = degrees of freedom. All models include Student and Language as random factors on the intercept.

| Combined Model classes | Combined model variables | dAICc | df |
|---|---|---|---|
| All four classes | B1_Urbanization + B2_Isolation^2 + B3_ParentEdu + B4_ParentJob^2 + D2_GradeEng^2 + D4_TradSkills + D5_ModSkills + C2_HomeLang^2 + C1_ParentMaxFluency + C3_SameLang + A1_LogN.2000. + A3_Endangered + A7_RegCode + A8_LogElevMedian | 0 | 23 |
| All four classes significant | B1_Urbanization + B2_Isolation^2 + B3_ParentEdu + B4_ParentJob^2 + D4_TradSkills + D5_ModSkills + C2_HomeLang^2 + C1_ParentMaxFluency + C3_SameLang + A3_Endangered + A7_RegCode | 0.14 | 19 |
| Socio-Economic + Student | B1_Urbanization + B2_Isolation^2 + B3_ParentEdu + B4_ParentJob^2 + D2_GradeEng^2 + D4_TradSkills + D5_ModSkills + C2_HomeLang^2 + C1_ParentMaxFluency + C3_SameLang | 48.56 | 17 |
| Socio-Economic + Family la | B1_Urbanization + B2_Isolation^2 + B3_ParentEdu + B4_ParentJob^2 + D2_GradeEng^2 + D4_TradSkills + D5_ModSkills + A1_LogN.2000. + A3_Endangered + A7_RegCode + A8_LogElevMedian | 146.59 | 19 |
| Socio-Economic + Family la | B1_Urbanization + B2_Isolation^2 + B3_ParentEdu + B4_ParentJob^2 + C2_HomeLang^2 + C1_ParentMaxFluency + C3_SameLang | 200.38 | 13 |
| Socio-Economic + Student | B1_Urbanization + B2_Isolation^2 + B3_ParentEdu + B4_ParentJob^2 + D2_GradeEng^2 + D4_TradSkills + D5_ModSkills | 223.06 | 15 |
| Student traits + Family lan | D2_GradeEng^2 + D4_TradSkills + D5_ModSkills + C2_HomeLang^2 + C1_ParentMaxFluency + C3_SameLang | 329.22 | 11 |
| Socio-Economic + Languag | B1_Urbanization + B2_Isolation^2 + B3_ParentEdu + B4_ParentJob^2 + A1_LogN.2000. + A3_Endangered + A7_RegCode + A8_LogElevMedian | 586.31 | 15 |
| Socio-Economic | B1_Urbanization + B2_Isolation^2 + B3_ParentEdu + B4_ParentJob^2 | 644.16 | 9 |
| Family language use | C2_HomeLang^2 + C1_ParentMaxFluency + C3_SameLang | 719.08 | 7 |
| Student traits | D2_GradeEng^2 + D4_TradSkills + D5_ModSkills C2_HomeLang^2 | 885.91 | 7 |
| Language traits | A1_LogN.2000. + A3_Endangered + A7_RegCode + A8_LogElevMedian | 1554.97 | 16 |
| Null | Null (random factors + intercept) | 1636.34 | 3 |

**Table S5.** Variables and their values used in the models extrapolating language fluency. The L1 language skills are extrapolated to the entire 18-20-year-old cohort in PNG (L1A model), to the 18-20-year-old cohort in PNG 30 years in the future (L1B), and to the current average socioeconomic parameters for the Lower-Middle Income countries (L1C). Language fluency (L2 = 3) is extrapolated to the to the entire 18-20-year-old cohort in PNG (L2A model), to the next generation of secondary students in PNG (L2B) and the next 18-20-year-old generation in PNG (L2C). See Materials and Methods for the procedure used to generate values for Adjusted variables.

| Model | Var# | Variable | Parameter values | Comments | Ref. | Data |
|---|---|---|---|---|---|---|
| L1A | A3 | LangStatus | 0 = 0.9782, 1 = 0.0218 | Distribution of PNG population among endangered/non-endangered languages based on Language size (A1) | (1) | External |
| L1A | A4 | LangRegion | 1 = 0.391, 2 = 0.279, 3 = 0.199, 4 = 0.131 | Distribution of PNG population among PNG regions, based on Language size (A1) | (1, 8) | External |
| L1A | B1 | Urbanization | 1 = 0.869, 3 = 0.131, 2 = not used | Current PNG urbanization rate | (2) | External |
| L1A | B2 | Remoteness | 1 = 0.772, 2 = 0.049, 3 = 0.036, 4 = 0.143 | Adjusted by Urbanization (B1) | | Adjusted |
| L1A | B3 | ParentEdu | 1 = 0.453, 2 = 0.354, 3 = 0.049, 4 = 0.076, 5 = 0.019, 6 = 0.049 | Education data for 50-54 years age group in PNG | (9) | External |
| L1A | B4 | ParentJob | 1 = 0.57, 2 = 0.125, 3 = 0.303 | Job structure in PNG 2009-2010 | (10) | External |
| L1A | C1 | ParentLang | 0 = 0.001, 1 = 0.009, 2 = 0.014, 3 = 0.976 | No change from original data | | This study |
| L1A | C2 | HomeLang | 1 = 0.469, 2 = 0.521, 3 = 0.010 | Adjusted by Urbanization (B1) & Parents' education (B3) | | Adjusted |
| L1A | C3 | SameLang | 0 = 0.240, 1 = 0.760 | Adjusted by Urbanization (B1) & Parents' education (B3) | | Adjusted |
| L1A | D2 | EnglishGrade | | Not used (part of population does not have Gr 10 test) | | Not used |
| L1A | D4 | TraditSkills | 0 = 0.010, 1 = 0.011, 2 = 0.041, 3 = 0.069, 4 = 0.106, 5 = 0.123, 6 = 0.154, 7 = 0.144, 8 = 0.162, 9 =0.117, 10 = 0.063 | Adjusted by Urbanization (B1) & Parents' education (B3) | | Adjusted |
| L1A | D5 | ModernSkills | 0 = 0.052, 1 = 0.203, 2 = 0.310, 3 = 0.316, 4 = 0.119 | Adjusted by Urbanization (B1) & Parents' education (B3) | | Adjusted |
| L1B | A3 | LangStatus | 0 = 0.922, 1 = 0.078 | Distribution of PNG population among endangered/non-endangered languages based on Language size (A1) | (1) | External |
| L1B | A4 | LangRegion | 1 = 0.391, 2 = 0.279, 3 = 0.199, 4 = 0.131 | Distribution of PNG population among PNG regions, based on Language size (A1) | (1, 8) | External |
| L1B | B1 | Urbanization | 1 = 0.76, 3 = 0.24, 2 = not used | Extrapolation for PNG in 2050 | (11) | External |
| L1B | B2 | Remoteness | 1 = 0.790, 2 = 0.045, 3 = 0.032, 4 = 0.133 | Adjusted by Urbanization (B1) | | Adjusted |
| L1B | B3 | ParentEdu | 1 = 0.254, 2 = 0.333, 3 = 0.100, 4 = 0.190, 5 = 0.057, 6 = 0.066 | Education of 30-34 years old | (9) | External |
| L1B | B4 | ParentJob | 1 = 0.57, 2 = 0.125, 3 = 0.303 | Data for PNG 2009-2010 were used as no trend was apparent over the past 3[illegible] | (10) | External |
| L1B | C1 | ParentLang | 0 = 0.0005, 1 = 0.010, 2 = 0.014, 3 = 0.976 | No change from original data | | This study |
| L1B | C2 | HomeLang | 0 = 0.329, 1 = 0.653, 3 = 0.018 | Adjusted by Family language uniformity (C3), Urbanization (B1) and Parents' education (B3) | | Adjusted |
| L1B | C3 | SameLang | 0 = 0.652, 1 = 0.348 | Estimated as the proportion of best friends speaking the same language (D7) | | This study |
| L1B | D2 | EnglishGrade | | Not used (part of population does not have Gr 10 test) | | This study |
| L1B | D4 | TraditSkills | 0 = 0.010, 1 = 0.013, 2 = 0.049, 3 = 0.082, 4 = 0.112, 5 = 0.132, 6 = 0.152, 7 = 0.142, 8 = 0.149, 9 =0.104, 10 = 0.055 | Adjusted by Urbanization (B1) & Parents' education (B3) | | Adjusted |
| L1B | D5 | ModernSkills | 0 = 0.038, 1 = 0.167, 2 = 0.287, 3 = 0.343, 4 = 0.165 | Adjusted by Urbanization (B1) & Parents' education (B3) | | Adjusted |
| L1C | A3 | LangStatus | 0 = 0.922, 1 = 0.078 | Distribution of PNG population among endangered/non-endangered languages based on Language size (A1) | (1) | External |
| L1C | A4 | LangRegion | 1 = 0.391, 2 = 0.279, 3 = 0.199, 4 = 0.131 | Distribution of PNG population among PNG regions, based on Language size (A1) | (1, 8) | External |
| L1C | B1 | Urbanization | 1 = 0.584, 2 = not used, 3 = 0.416 | Lower-middle income countries in 2020 | (11) | External |
| L1C | B2 | Remoteness | 1 = 0.818, 2 = 0.038, 3 = 0.028, 4 = 0.116 | Adjusted by Urbanization (B1) | | This study |
| L1C | B3 | ParentEdu | 1 = 0.247, 2.5 = 0.430, 4.5 = 0.276, 6 = 0.047 | Education attainment in 30-40yrs old population in lower-middle income countries | (12) | External |
| L1C | B4 | ParentJob | 1.5 = 0.39, 3 = 0.61 | 1.5 = work in agriculture, 3 = employment in other sectors | (12) | External |
| L1C | C1 | ParentLang | 0 = 0.0005, 1 = 0.010, 2 = 0.014, 3 = 0.976 | No change from original data | | This study |
| L1C | C2 | HomeLang | 0 = 0.269, 1 = 0.702, 3 = 0.029 | Adjusted by Family language uniformity (C3), Urbanization (B1) and Parents' education (B3) | | Adjusted |
| L1C | C3 | SameLang | 0 = 0.652, 1 = 0.348 | Estimated as the proportion of best friends speaking the same language (D7) | | This study |
| L1C | D4 | TraditSkills | 0 = 0.010, 1 = 0.016, 2 = 0.056, 3 = 0.097, 4 = 0.117, 5 = 0.135, 6 = 0.151, 7 = 0.130, 8 = 0.144, 9 =0.093, 10 = 0.051 | Adjusted by Urbanization (B1) & Parents' education (B3) | | Adjusted |
| L1C | D5 | ModernSkills | 0 = 0.049, 1 = 0, 2 = 0.608, 3 = 0.142, 4 = 0.201 | Based on access to mobile phone and computer and access to internet | (12) | External |

**Table S5.** Continued

| Model | Var# | Variable | Parameter values | Comments | Ref. | Data |
|---|---|---|---|---|---|---|
| L2A | B1 | Urbanization | 1 = 0.869, 3 = 0.131, 2 = not used | Current urbanization rate | (2) | External |
| L2A | B3 | ParentEdu | 1 = 0.453, 2 = 0.354, 3-6 = 0.193 | Education data for 50-54 years age group in PNG | (9) | External |
| L2A | B4 | ParentJob | 1-2 = 0.695, 3 = 0.305 | Job structure in PNG 2009-2010 | (10) | External |
| L2A | C2 | HomeLang | 1 = 0.467 2-3 = 0.533 | Adjusted by Urbanization (B1) & Parents' education (B3) | | Adjusted |
| L2A | C3 | SameLang | 0 = 0.244, 1 = 0.756 | Adjusted by Urbanization (B1) & Parents' education (B3) | | Adjusted |
| L2B | B1 | Urbanization | 1 = 0.76, 3 = 0.24, 2 = not used | Extrapolation for PNG in 2050 | (11) | External |
| L2B | B3 | ParentEdu | 1 = 0, 2 = 0, 3 0, 4 = 0, 5 = 0.532, 6 = 0.468 | Proportion of secondary school graduates with tertiary education for 25-34 age group in PNG | (9) | External |
| L2B | C2 | HomeLang | 1 = 0.306, 2-3 = 0.694 | Adjusted by Family language unifornity (C3), Urbanization (B1), Parents' education (B3), and Parent's Language skills (C1) (vernacular language use only for those fluent in it, L2 = 3) | | Adjusted |
| L2B | C3 | SameLang | 0 = 0.556, 1 = 0.444 | The proportion of best friends speaking the same language for those fluent in language (D7) | | This study |
| L2C | B1 | Urbanization | 1 = 0.76, 3 = 0.24, 2 = not used | Extrapolation for PNG in 2050 | (11) | External |
| L2C | B3 | ParentEdu | 1 = 0.254, 2 = 0.333, 3 = 0.100, 4 = 0.190, 5 = 0.057, 6 = 0.066 | Education of 30-34 years old in PNG | (9) | External |
| L2C | B4 | ParentJob | 1 = 0.572, 2 = 0.125, 3 = 0.303 | Job structure in PNG 2009-2010 used as no trends in PNG jobs apparent | (11) | External |
| L2C | C2 | HomeLang | 1 = 0.388 2-3 = 0.612 | Adjusted by Urbanization (B1), Parents' education (B3) and Parent's language skills (C1) (vernacular language use only for those fluent in it, L2 = 3) | | Adjusted |
| L2C | C3 | SameLang | 0 = 0.297, 1 = 0.703 | Adjusted by Urbanization (B1), Parents' education (B3) and Parent's language skills (C1) (vernacular language use only for those fluent in it, L2 = 3) | | Adjusted |

**Table S6.** Model selection results for the candidate models assessing the role of socioeconomic variables B1 – B4, family language use variables C1 – C3, and student-related variables D1 – D6 on the knowledge of bird species (E1) and traditional plant use (E2) by students. dAICc = delta corrected Akaike Information Criterion, df = degrees of freedom. All models include Student and Language as random factors on the intercept.

| Taxon | Variable class | Models – Student trait variables | dAICc | df |
|---|---|---|---|---|
| Birds | Socio-economic | B1_Urbanization + B2_Isolation^2 + B3_ParentEdu^2 + B4_ParentJob^2 | 0 | 10 |
| Birds | Socio-economic | B1_Urbanization + B2_Isolation^3 + B3_ParentEdu + B4_ParentJob^2 | 4.60 | 10 |
| Birds | Socio-economic | B1_Urbanization + B2_Isolation^2 + B3_ParentEdu + B4_ParentJob^2 | 4.80 | 9 |
| Birds | Socio-economic | B1_Urbanization^2 + B2_Isolation^3 + B3_ParentEdu + B4_ParentJob^2 | 4.89 | 11 |
| Birds | Socio-economic | B1_Urbanization + B2_Isolation^2 + B3_ParentEdu^2 + B4_ParentJob | 7.63 | 9 |
| Birds | Socio-economic | B1_Urbanization + B2_Isolation^2 + B3_ParentEdu | 17.15 | 7 |
| Birds | Socio-economic | B1_Urbanization + B2_Isolation + B3_ParentEdu + B4_ParentJob^2 | 28.68 | 8 |
| Birds | Socio-economic | B1_Urbanization + B3_ParentEdu + B4_ParentJob^2 | 51.05 | 7 |
| Birds | Socio-economic | B1_Urbanization + B2_Isolation^2 + B4_ParentJob^2 | 90.28 | 8 |
| Birds | Socio-economic | B2_Isolation^2 + B3_ParentEdu + B4_ParentJob^2 | 357.64 | 8 |
| Birds | Socio-economic | Null (random factors + intercept) | 835.38 | 3 |
| Birds | Language use | C2_HomeLang^2 + C1_ParentMaxFluency^2 + C3_SameLang | 0 | 8 |
| Birds | Language use | C2_HomeLang + C1_ParentMaxFluency + C3_SameLang | 2.54 | 6 |
| Birds | Language use | C2_HomeLang^2 + C1_ParentMaxFluency + C3_SameLang | 6.56 | 7 |
| Birds | Language use | C2_HomeLang + C3_SameLang | 8.43 | 5 |
| Birds | Language use | C2_HomeLang, 2 + C1_ParentMaxFluency, 1 | 132.04 | 6 |
| Birds | Language use | C3_SameLang | 349.94 | 4 |
| Birds | Language use | Null (random factors + intercept) | 569.16 | 3 |
| Birds | Student traits | D1_Gender + D2_GradeEng^3 + D4_TradSkills + D5_ModSkills | 0 | 9 |
| Birds | Student traits | D1_Gender + D2_GradeEng^2 + D4_TradSkills + D5_ModSkills | 1.99 | 8 |
| Birds | Student traits | D1_Gender + D2_GradeEng^3 + D3_GradeMath^3 + D4_TradSkills + D5_ModSkills | 5.17 | 12 |
| Birds | Student traits | D1_Gender + D2_GradeEng + D4_TradSkills + D5_ModSkills | 19.08 | 7 |
| Birds | Student traits | D1_Gender + D2_GradeEng^3 + D3_GradeMath^3 + D6_Trad.Mod | 22.33 | 11 |
| Birds | Student traits | D1_Gender + D4_TradSkills + D5_ModSkills | 42.13 | 6 |
| Birds | Student traits | D2_GradeEng^3 + D4_TradSkills + D5_ModSkills | 81.35 | 8 |
| Birds | Student traits | D2_GradeEng^3 + D3_GradeMath^3 + D4_TradSkills + D5_ModSkills | 81.96 | 11 |
| Birds | Student traits | D2_GradeEng^2 + D4_TradSkills + D5_ModSkills | 84.53 | 7 |
| Birds | Student traits | D2_GradeEng + D4_TradSkills + D5_ModSkills | 100.34 | 6 |
| Birds | Student traits | D4_TradSkills + D5_ModSkills | 125.79 | 5 |
| Birds | Student traits | D2_GradeEng^3 + D6_Trad.Mod | 162.11 | 7 |
| Birds | Student traits | D1_Gender + D2_GradeEng^3 + D3_GradeMath^3 + D4_TradSkills | 180.34 | 11 |
| Birds | Student traits | D1_Gender + D2_GradeEng^3 + D3_GradeMath^3 + D5_ModSkills | 243.15 | 11 |
| Birds | Student traits | Null (random factors + intercept) | 1066.42 | 3 |

**Table S6.** Continued.

| Taxon | Variable class | Models – Student trait variables | dAICc | df |
|---|---|---|---|---|
| Plants | Socio-economic | B1_Urbanization + B2_Isolation^2 + B3_ParentEdu + B4_ParentJob | 0 | 8 |
| Plants | Socio-economic | B1_Urbanization + B2_Isolation^2 + B3_ParentEdu + B4_ParentJob^2 | 0.33 | 9 |
| Plants | Socio-economic | B1_Urbanization + B2_Isolation^2 + B3_ParentEdu^2 + B4_ParentJob | 1.49 | 9 |
| Plants | Socio-economic | B1_Urbanization + B2_Isolation^2 + B3_ParentEdu^2 + B4_ParentJob^2 | 1.95 | 10 |
| Plants | Socio-economic | B1_Urbanization + B2_Isolation^3 + B3_ParentEdu + B4_ParentJob^2 | 2.27 | 10 |
| Plants | Socio-economic | B1_Urbanization^2 + B2_Isolation^3 + B3_ParentEdu + B4_ParentJob^2 | 4.27 | 11 |
| Plants | Socio-economic | B1_Urbanization + B2_Isolation^2 + B3_ParentEdu | 5.12 | 7 |
| Plants | Socio-economic | B1_Urbanization + B2_Isolation + B3_ParentEdu + B4_ParentJob | 14.16 | 7 |
| Plants | Socio-economic | B1_Urbanization + B2_Isolation + B3_ParentEdu + B4_ParentJob^2 | 14.22 | 8 |
| Plants | Socio-economic | B1_Urbanization + B3_ParentEdu + B4_ParentJob^2 | 24.50 | 7 |
| Plants | Socio-economic | B1_Urbanization + B2_Isolation^2 + B4_ParentJob^2 | 25.19 | 8 |
| Plants | Socio-economic | B1_Urbanization + B2_Isolation^2 + B4_ParentJob | 27.28 | 7 |
| Plants | Socio-economic | B2_Isolation^2 + B3_ParentEdu + B4_ParentJob^2 | 181.05 | 8 |
| Plants | Socio-economic | Null (random factors + intercept) | 392.61 | 3 |
| Plants | Language use | C2_HomeLang^2 + C1_ParentMaxFluency + C3_SameLang | 0 | 7 |
| Plants | Language use | C2_HomeLang + C3_SameLang | 0.57 | 5 |
| Plants | Language use | C2_HomeLang^2 + C1_ParentMaxFluency^2 + C3_SameLang | 1.70 | 8 |
| Plants | Language use | C2_HomeLang + C1_ParentMaxFluency + C3_SameLang | 1.72 | 6 |
| Plants | Language use | C2_HomeLang^2 + C1_ParentMaxFluency | 37.32 | 6 |
| Plants | Language use | +C3_SameLang | 185.97 | 4 |
| Plants | Language use | Null (random factors + intercept) | 263.85 | 3 |
| Plants | Student traits | D1_Gender + D2_GradeEng^2 + D4_TradSkills + D5_ModSkills | 0 | 8 |
| Plants | Student traits | D1_Gender + D2_GradeEng^3 + D4_TradSkills + D5_ModSkills | 0.25 | 9 |
| Plants | Student traits | D1_Gender + D2_GradeEng^3 + D3_GradeMath^3 + D4_TradSkills + D5_ModSkills | 1.79 | 12 |
| Plants | Student traits | D1_Gender + D2_GradeEng^3 + D3_GradeMath^3 + D6_Trad.Mod | 2.63 | 11 |
| Plants | Student traits | D2_GradeEng^2 + D4_TradSkills + D5_ModSkills | 5.28 | 7 |
| Plants | Student traits | D2_GradeEng^3 + D4_TradSkills + D5_ModSkills | 5.31 | 8 |
| Plants | Student traits | D2_GradeEng^3 + D3_GradeMath^3 + D4_TradSkills + D5_ModSkills | 6.66 | 11 |
| Plants | Student traits | D1_Gender + D2_GradeEng + D4_TradSkills + D5_ModSkills | 6.70 | 7 |
| Plants | Student traits | D1_Gender + D4_TradSkills + D5_ModSkills | 11.44 | 6 |
| Plants | Student traits | D2_GradeEng + D4_TradSkills + D5_ModSkills | 11.68 | 6 |
| Plants | Student traits | D2_GradeEng^3 + D6_Trad.Mod | 13.18 | 7 |
| Plants | Student traits | D4_TradSkills + D5_ModSkills | 16.85 | 5 |
| Plants | Student traits | D1_Gender + D2_GradeEng^3 + D3_GradeMath^3 + D5_ModSkills | 94.18 | 11 |
| Plants | Student traits | D1_Gender + D2_GradeEng^2 + D4_TradSkills | 94.81 | 7 |
| Plants | Student traits | D1_Gender + D2_GradeEng^3 + D3_GradeMath^3 + D4_TradSkills | 96.51 | 11 |
| Plants | Student traits | Null (random factors + intercept) | 369.48 | 3 |

**Table S7.** Model selection results for the candidate models assessing the combined role of the four variable classes A – D on the knowledge of bird species (E1) and traditional plant use (E2) by students. dAICc = delta corrected Akaike Information Criterion, df = degrees of freedom. All models include Student and Language as random factors on the intercept.

| Taxon | Combined Model classes | Combined model variables | dAICc | df |
|---|---|---|---|---|
| Birds | Socio-Economic + Student | L1_BodyParts^2 + B1_Urbanization + B3_ParentEdu^2 + D1_Gender + D4_TradSkills + D5_ModSkills | 0 | 11 |
| Birds | Socio-Economic + Student | L1_BodyParts^2 + B1_Urbanization + B2_Isolation^2 + B3_ParentEdu^2 + B4_ParentJob^2 + D1_Gender + D2_GradeEng^2 + D4_TradSkills + D5_ModSkills | 3.30 | 17 |
| Birds | Socio-Economic + Student | L1_BodyParts^2 + B1_Urbanization + B2_Isolation^2 + B3_ParentEdu^2 + B4_ParentJob^2 + D1_Gender + D2_GradeEng^2 + D4_TradSkills + D5_ModSkills + C2_HomeLang^2 + C1_ParentMaxFluency^2 + C3_SameLang | 4.10 | 22 |
| Birds | Student traits + Family lan | L1_BodyParts^2 + D1_Gender + D2_GradeEng^2 + D4_TradSkills + D5_ModSkills + C2_HomeLang^2 + C1_ParentMaxFluency^2 + C3_SameLang | 44.70 | 15 |
| Birds | Socio-Economic + Family la | L1_BodyParts^2 + B1_Urbanization + B2_Isolation^2 + B3_ParentEdu^2 + B4_ParentJob^2 + C2_HomeLang^2 + C1_ParentMaxFluency^2 + C3_SameLang | 387.30 | 17 |
| Birds | Language skills | L1_BodyParts^2 | 572.70 | 5 |
| Birds | Socio-Economic + Student | B1_Urbanization + B2_Isolation^2 + B3_ParentEdu^2 + B4_ParentJob^2 + D1_Gender + D2_GradeEng^2 + D4_TradSkills + D5_ModSkills | 1467.70 | 15 |
| Birds | Student traits | D1_Gender + D2_GradeEng^2 + D4_TradSkills + D5_ModSkills | 1799.80 | 9 |
| Birds | Socio-Economic | B1_Urbanization + B2_Isolation^2 + B3_ParentEdu^2 + B4_ParentJob^2 | 2038.40 | 9 |
| Birds | Family language use | C2_HomeLang^2 + C1_ParentMaxFluency^2 + C3_SameLang | 2299.60 | 6 |
| Birds | Null | Null (random factors + intercept) | 2866.20 | 3 |
| Plants | Socio-Economic + Student | L1_BodyParts^2 + B1_Urbanization + D1_Gender + D4_TradSkills + D5_ModSkills + C2_HomeLang | 0 | 10 |
| Plants | Socio-Economic + Student | L1_BodyParts^2 + B1_Urbanization + B2_Isolation^2 + B3_ParentEdu + B4_ParentJob + D1_Gender + D2_GradeEng^2 + D4_TradSkills + D5_ModSkills + C2_HomeLang + C3_SameLang | 5.20 | 17 |
| Plants | Socio-Economic + Student | L1_BodyParts^2 + B1_Urbanization + B2_Isolation^2 + B3_ParentEdu + B4_ParentJob + D1_Gender + D2_GradeEng^2 + D4_TradSkills + D5_ModSkills | 10.10 | 15 |
| Plants | Student traits + Family lan | L1_BodyParts^2 + B1_Urbanization + B2_Isolation^2 + B3_ParentEdu + B4_ParentJob + D1_Gender + D2_GradeEng^2 + D4_TradSkills + D5_ModSkills + C2_HomeLang + C3_SameLang | 28.00 | 12 |
| Plants | Socio-Economic + Family la | L1_BodyParts^2 + B1_Urbanization + B2_Isolation^2 + B3_ParentEdu + B4_ParentJob + D1_Gender + D2_GradeEng^2 + D4_TradSkills + D5_ModSkills + C2_HomeLang + C3_SameLang | 62.00 | 12 |
| Plants | Socio-Economic + Student | L1_BodyParts^2 + B1_Urbanization + B2_Isolation^2 + B3_ParentEdu + B4_ParentJob + D1_Gender + D2_GradeEng^2 + D4_TradSkills + D5_ModSkills + C2_HomeLang + C3_SameLang | 408.60 | 15 |
| Plants | Socio-Economic | L1_BodyParts^2 + B1_Urbanization + B2_Isolation^2 + B3_ParentEdu + B4_ParentJob + D1_Gender + D2_GradeEng^2 + D4_TradSkills + D5_ModSkills + C2_HomeLang + C3_SameLang | 601.00 | 9 |
| Plants | Student traits | L1_BodyParts^2 + B1_Urbanization + B2_Isolation^2 + B3_ParentEdu + B4_ParentJob + D1_Gender + D2_GradeEng^2 + D4_TradSkills + D5_ModSkills + C2_HomeLang + C3_SameLang | 622.90 | 9 |
| Plants | Family language use | L1_BodyParts^2 + B1_Urbanization + B2_Isolation^2 + B3_ParentEdu + B4_ParentJob + D1_Gender + D2_GradeEng^2 + D4_TradSkills + D5_ModSkills + C2_HomeLang + C3_SameLang | 730.00 | 6 |
| Plants | Language skills | L1_BodyParts^2 + B1_Urbanization + B2_Isolation^2 + B3_ParentEdu + B4_ParentJob + D1_Gender + D2_GradeEng^2 + D4_TradSkills + D5_ModSkills + C2_HomeLang + C3_SameLang | 861.50 | 4 |
| Plants | Null | Null (random factors + intercept) | 992.10 | 3 |

**SI References**

1. D. M. Eberhard, G. F., Simons, C. D. Fennig, Eds., *Ethnologue: Languages of the World* (SIL International, ed. 23, 2020).
2. Worldometers, *Papua New Guinea Population* (March 2020); https://www.worldometers.info/world-population/papua-new-guinea-population/.
3. National Statistical Office (NSO), *Papua New Guinea 2000 Census: Final Figures* (NSO, Port Moresby, 2002).
4. A. Jarvis, H. I. Reuter, A. Nelson, E. Guevara, Hole-filled seamless SRTM data V4. (Int. Ctr. Trop. Agric. CIAT (September 2020); http://srtm.csi.cgiar.org.
5. PNG Bible Translation Association (PNGBTA), *Papua New Guinea Scriptures* (March 2020); https://png.bible/.
6. S. J. Greenhill, *TransNewGuinea.org, a Database of the Languages of New Guinea* March 2020); http://transnewguinea.org/.
7. R. Cámara–Leret, Z. Dennehy, Indigenous knowledge of New Guinea's useful plants: a review, *Econ. Bot.* **73**, 405–415 (2019).
8. H. Hammarström, R. Forkel, M. Haspelmath, S. Bank, Sebastian, *Glottolog 4.2.1*. (Max Planck Inst. Science of Human History, Jena (March 2020); http://glottolog.org.
9. National Statistical Office (NSO), *Papua New Guinea Demographic and Health Survey 2016-18* (NSO, 2019).
10. National Statistical Office (NSO). *2009-2010 Papua New Guinea Household Income and Expenditure Survey*, (NSO, 2011).
11. United Nations, *World Urbanization Prospects: The 2018 Revision*, (United Nations, 2019).
12. World Bank, *Data Catalog: Educational Statistics* (March 2020); https://datacatalog.worldbank.org/dataset/education-statistics.
